# BRCA2 loss triggers a downward spiral of genomic instability via ROS-dependent metabolic collapse

**DOI:** 10.64898/2026.05.15.725601

**Authors:** Hayato Hirai, Tomohiro Iguchi, Kenji Kitajima, Kosuke Yamazaki, Daichi Sadato, Kosuke Tanegashima, Yutaka Kanoh, Hiroko Masuda, Saori Wakasa, Manami Sano, Ryotaro Kawasumi, Kosei Yamada, Shintaro Yamada, Akinori Endo, Noriko Nakatsugawa, Kazuto Takayasu, Novokreshchenov Leonid, Midori Yamaguchi, Hiroshi Shitara, Ken-ichiro Hayashi, Takuma Kumamoto, Yasumasa Nishito, Ryo Shioda, Ngo Thi To Trinh, Toshiyasu Taniguchi, Shinichiro Nakada, Masato T. Kanemaki, Ichiro Taniuchi, Hideya Kawaji, Junko Murai, Kouji Hirota, Kousuke Yusa, Masakazu Toi, Hiroyuki Sasanuma

**Affiliations:** Department of Genome Dynamics, Tokyo Metropolitan Institute of Medical Science, Tokyo, 156-8506, Japan; Department of Computational Biology and Medical Sciences, Graduate School of Frontier Sciences, The University of Tokyo, 5-1-5 Kashiwanoha, Kashiwa-shi, Chiba, 277-8563, Japan; Clinical Research and Trials Center, Tokyo Metropolitan Komagome Hospital, 3-18-22 Honkomagome, Bunkyo-ku, Tokyo, 113-8677, Japan; Department of Breast Surgery, Tokyo Metropolitan Cancer and Infectious Diseases Center Komagome Hospital, 3-18-22 Honkomagome, Bunkyo-ku, Tokyo, 113-8677, Japan; Department of Chemistry, Graduate School of Science, Tokyo Metropolitan University, Minamiosawa 1-1, Hachioji-shi, Tokyo 192-0397, Japan; Faculty of Medicine, Kyoto University, Yoshida-Konoe-cho, Sakyo-ku, Kyoto 606-8501, Japan; Laboratory of Protein Metabolism, Tokyo Metropolitan Institute of Medical Science, Tokyo, 156-8506, Japan; Animal Research Division, Center for Basic Technology Research, Tokyo Metropolitan Institute of Medical Science, Tokyo, 156-8506, Japan; Department of Bioscience, Okayama University of Science, 1-1 Ridai-cho, Okayama 700-0005, Japan; Developmental Neuroscience Project, Tokyo Metropolitan Institute of Medical Science, Tokyo, 156-8506, Japan; Technology Research Division, Center for Basic Technology Research, Tokyo Metropolitan Institute of Medical Science, Tokyo, 156-8506, Japan; Revvity Japan K.K., 134 Godo-cho, Hodogaya-ku, Yokohama, Kanagawa 240-0005, Japan; Department of Molecular Life Science, Tokai University School of Medicine, 143 Shimokasuya, Isehara, Kanagawa, 259-1193, Japan; Department of Biochemistry and Molecular Biology, Graduate School of Medical Science, Kyoto Prefectural University of Medicine, Kyoto, 602-8566, Japan; Department of Chromosome Science, National Institute of Genetics, Research Organization of Information and Systems (ROIS), Mishima, Shizuoka, 411-8540, Japan; Graduate Institute for Advanced Studies, SOKENDAI, Yata 1111, Mishima, Shizuoka 411-8540, Japan; Department of Biological Science, Graduate School of Science, The University of Tokyo, Bunkyo-ku, Tokyo 113-0033, Japan; Laboratory for Transcriptional Regulation, RIKEN Center for Integrative Medical Sciences (IMS), Yokohama, Kanagawa, 230-0045, Japan; Research Center for Genome & Medical Sciences, Tokyo Metropolitan Institute of Medical Science, 2-1-6 Kamikitazawa, Setagaya-ku, Tokyo, 156-8506, Japan; Department of Cell Growth and Tumor Regulation, Proteo-Science Center, Premier Institute for Advanced Studies, Ehime University, Toon, Ehime, 791-0295, Japan; Department of Biochemistry and Molecular Genetics, Graduate School of Medicine, Ehime University, Toon, Ehime, 791-0295, Japan; Department of Stem Cell Genetics, Institute for Life and Medical Sciences, Kyoto University, 53 Shogoin Kawahara-cho, Sakyo-ku, Kyoto, 606-8507, Japan; Tokyo Metropolitan Cancer and Infectious Disease Center, Komagome Hospital, Tokyo, 113-8677, Japan

**Keywords:** BRCA2, Reactive oxygen species, Homologous recombination

## Abstract

BRCA2 plays a central role in maintaining genome integrity through homologous recombination and replication-fork protection, yet the compensatory networks sustaining *BRCA2*-deficient cells remain unclear. Here we show BRCA2 enforces a homeostatic mechanism aligning mitochondrial respiration with DNA repair capacity in both cancer and non-cancer contexts. Genome-wide CRISPR screening identified glutathione metabolism and base-excision repair as the key compensatory networks sustaining *BRCA2*-deficient cells by detoxifying mitochondria-derived reactive oxygen species. BRCA2 loss provokes an acute mitochondrial ROS surge, causing 8-oxoguanine accumulation and a systemic metabolic crisis marked by NAD^+^ and glutathione depletion. PARP inhibitor targets DNA replication vulnerabilities, increasing the cellular requirement for BRCA2. The resulting oxidative burden primes cells for *TP53*-dependent apoptosis in G_1_ during olaparib treatment, which extends cytotoxicity beyond canonical S-phase stress. These findings indicate BRCA2 prevents metabolic flux from outpacing repair capacity, providing a rationale for combining PARP inhibition with redox modulation to enhance efficacy and overcome resistance.

**Highlights:**

- Acute BRCA2 loss induces ROS and mitochondrial dysfunction creating a metabolic scar
- Oxidative lesions drive PARP hyperactivation and precipitate a cellular NAD crisis
- PARP inhibitors provoke TP53-dependent apoptosis in G_1_ beyond replication stress in S phase
- Glutathione deficiency exacerbates bone marrow failure under BRCA2 depletion
- BRCA2 tightly couples mitochondrial redox homeostasis to genomic maintenance

## Introduction

Cells must continually balance the demands of energy production with the rigorous maintenance of genomic integrity^1,2^. Mitochondrial metabolism, while indispensable for bioenergetics, inevitably generates reactive oxygen species (ROS) as byproducts, posing a persistent threat to the genome^3^. In response, cells activate sophisticated antioxidant programs, such as the Keap1-Nrf2 axis, to maintain redox homeostasis^4,5^. To further counteract these pervasive threats and resolve oxidative DNA lesions, cells are thought to recruit a robust network of tumor suppressors and DNA repair factors. Among these, BRCA2, as well as BRCA1, is established as a central regulator for maintaining genomic integrity. However, the precise mechanisms by which these regulators coordinate cellular responses to continuous oxidative stress, and the compensatory pathways that sustain cellular homeostasis upon their loss, remain largely unknown.

BRCA2 protein is essential for repairing double-strand breaks via homologous recombination (HR) and protecting stalled replication forks^6,7^. Consequently, their loss underlies hereditary breast and ovarian cancer (HBOC) and confers an exquisite sensitivity to PARP inhibitors (PARPi)^8–11^. Numerous studies have attributed this PARPi lethality primarily to S-phase-specific mechanisms, such as replication stress and unresolved damage at replication forks^11–13^. Physiological ROS target guanine to generate 8-oxo-7,8-dihydroguanine (8-oxoG), a highly mutagenic oxidative DNA lesion^14,15^. These lesions are primarily processed via the base excision repair (BER) pathway, a process that inherently generates hazardous single-strand break (SSB) intermediates. If replication forks encounter these intermediates during BER, it can trigger fork collapse, transforming the repair process itself into a primary driver of genomic instability. Previous reports have established that *BRCA*-deficient cells exhibit a critical dependency on BER factors, such as Polβ, Apex2 and Fen1, for survival^16–18^. However, the fundamental basis of this dependency on the BER pathway and how BRCA2 deficiency reshapes global redox and metabolic homeostasis to create vulnerabilities beyond S-phase remain unexplored.

In this study, we demonstrated that BRCA2 acts as a key regulator of redox-metabolic networks to maintain cellular homeostasis. BRCA2 loss triggered mitochondrial hyperactivation and a ROS surge, leading to 8-oxoG accumulation in open chromatin, PARP hyperactivation, and the exhaustion of NAD^+^ and glutathione pools. This systemic metabolic collapse precipitated *TP53*-dependent apoptosis in G_1_, expanding the mechanism of PARPi lethality beyond canonical S-phase stress.

Furthermore, mouse models revealed this redox-metabolic derangement as the primary driver of PARPi-associated bone marrow toxicity. Our findings establish that BRCA2 ensures cellular metabolic activity does not outpace repair capacity, providing a compelling rationale for synergistic PARPi and redox therapies.

## Result

### An unbiased CRISPR screen maps compensatory networks engaged by BRCA2 depletion

The cellular mechanisms that compensate for BRCA2 loss remain poorly understood. To address this challenge directly, we set out to systematically and comprehensively define the genetic pathways that sustain cell fitness following acute BRCA2 depletion. We therefore performed a genome-wide CRISPR/Cas9 loss-of-function screen in *TP53*-proficient, near-diploid TK6 cells, using an auxin-inducible degron (AID) system to achieve acute BRCA2 depletion^19,20^(**Figures 1A, S1A, and S1B**). BRCA2^AID^ cells were generated by C-terminal mAID tagging (**Figure S1A**). Auxin treatment efficiently depleted BRCA2, resulted in slower growth, and reduced RAD51 foci, validating the functionality of this system (**Figures 1B, S1C, and S1D**). Following library transduction, we contrasted guide RNA enrichment between auxin-treated and control populations to identify modifiers of *BRCA2*-deficient fitness (**Figure 1C and Table S1-3**). To validate synthetic-lethal dependencies, we individually disrupted 18 top-ranked candidates. Among these, 16 genes, including PARP1, significantly reduced proliferation specifically under BRCA2 depletion (**Figure S1E**). Notably, the screen also identified *TP53* as a primary suppressor of growth inhibition induced by BRCA2 loss (**Figure S1F**). We confirmed this by generating *BRCA2*^AID^ *TP53*^−/−^ cells, where *TP53* deficiency rescued cell proliferation and restored survival to near-*wild-type* levels upon BRCA2 depletion (**Figures S1G and S1H**). Collectively, these results provide an unbiased map of the pathways sustaining *BRCA2*-deficient cells and establish a robust platform for subsequent mechanistic analyses.

**Figure 1.**
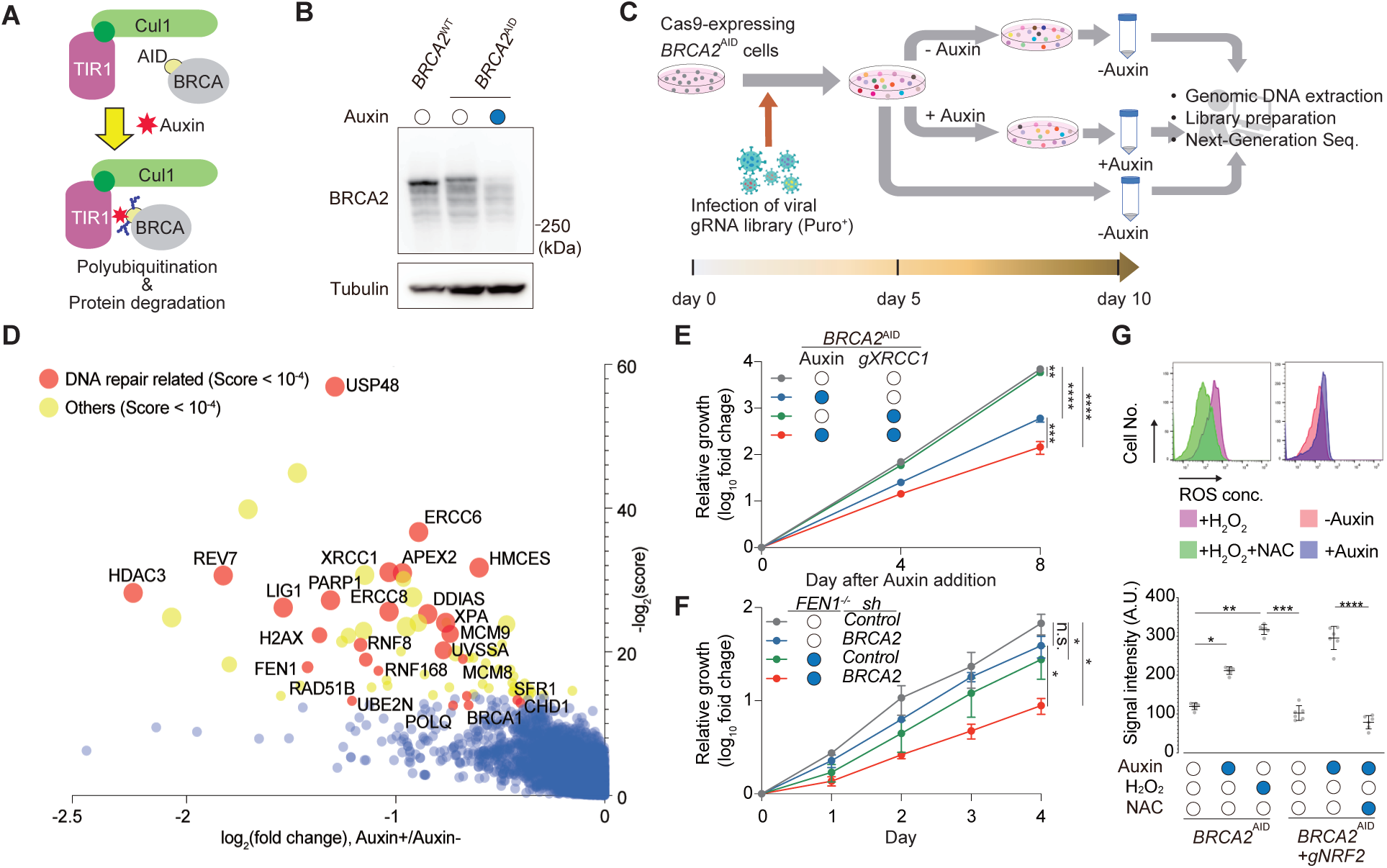
BRCA2 depletion induces dependency on oxidative DNA damage repair pathways. (A) Schematic of the Auxin-Inducible Degron (AID) system. In the presence of OsTIR1, the addition of auxin (IAA) mediates the ubiquitylation and proteasomal degradation of mAID-tagged BRCA2. (B) Immunoblot showing time-dependent degradation of BRCA2 upon 30 µM auxin treatment. Tubulin serves as a loading control. (C) Experimental workflow for the whole-genome CRISPR/Cas9 knockout screen. Cas9-expressing *BRCA2*^AID^ cells were infected with a viral gRNA library and cultured with or without 30 µM auxin to identify genetic interactions with BRCA2 loss. (D) Volcano plot displaying CRISPR screening results. The x-axis represents the fold change (Auxin^+^/Auxin^−^) with log_2_, and the y-axis represents the significance (-log_2_(score)). Red circles indicate DNA repair genes with scores < 10^−4^. (E and F) Validation of the screening hits. Growth curves of *BRCA2*^AID^ cells transduced with sgRNAs targeting *XRCC1* gene in the presence or absence of 500µM auxin (E). Growth curves of *FEN1*-proficient and *FEN1*-deficient cells transduced with shRNAs targeting *BRCA2* (F). (G) Intracellular ROS levels analyzed by flow cytometer. BRCA2 depletion (500 µM auxin, 24 h) increases ROS, which is rescued by 1 mM N-acetylcysteine (NAC). Data are presented as mean ± SD from at least three-independent experiments. Statistical significance was calculated using an unpaired t-test (E and F) or Mann-Whitney test (G). \**p* < 0.05, \*\**p* < 0.01, \*\*\**p* < 0.001, \*\*\*\**p*<0.0001.

### BRCA2 deficiency confers synthetic lethality with the base excision repair (BER) pathway

Our results revealed a broad spectrum of mutations affecting diverse DNA repair and DNA damage response pathways **(Figure 1D)**, similar to those observed in previous CRISPR-based screens using *BRCA2*-deficient cancer cells^16,21^. For example, we identified components of HR (e.g., *RAD51B*, *SFR1*, and *MCM9*)^22–25^, ubiquitin-mediated signaling factors involved in DNA repair (*e.g*., *RNF8, RNF168, UBE2N,* and *USP48*)^26–28^, and additional factors participating in alternative repair mechanisms, including *POLQ* for microhomology-mediated end joining^29^, *REV7* for translesion synthesis, and *XPA*, *ERCC6*, and *UVSSA* for NER/TC-NER pathways^30^. Additionally, as demonstrated before^16,21^, we also identified multiple base excision repair (BER) factors, *e.g.*, *PARP1*, *XRCC1*, *APEX2*, *FEN1*, and *LIG1*^31,32^ (**Figure 1D)**. To validate the synthetic defect between BRCA2 loss and BER mutants, we analyzed the effect of *XRCC1* deletion in *BRCA2*^AID^ cells. We found that BRCA2-depleted cells lacking *XRCC1* not only exhibited slower proliferation compared with control cells but also showed increased sensitivity to hydrogen peroxide, H₂O₂ (**Figures 1E, S1I, and S1J**). Furthermore, the depletion of either BRCA1 or BRCA2 using *shRNA* in *FEN1^−/−^* cells significantly reduced cell proliferation, compared with single *FEN1^−/−^*cells^31^ (**Figures 1F, S1K, and S1L**). These findings indicate that BRCA2 depletion, possibly also BRCA1 depletion, markedly increases cellular dependency on the BER pathway, highlighting that deficiency of BER process in this context can lead to synthetic lethality.

### BRCA2 deficiency increases ROS

The BER pathway repairs DNA base lesions caused by oxidative stress, such as those induced by ROS. We hypothesized that BRCA2 depletion promotes a pro-oxidative state, increasing reliance on BER for genome integrity. To test this, we measured intracellular ROS after 24 h of auxin treatment using a fluorescent probe in TK6 cells (**Figure 1G**). BRCA2-depleted cells showed significant ROS accumulation compared to controls (**Figure 1G**), indicating elevated oxidative stress upon BRCA2 loss. ROS is sensed by the KEAP1-NRF2 axis^5^, which activates antioxidant gene expression^33^. To assess its role, we deleted *NRF2* in *BRCA2*^AID^ cells (**Figure S1I**). Combined loss of BRCA2 and NRF2 further increased ROS, which was reversed by NAC (N-Acetylcysteine), confirming NRF2’s critical function in limiting oxidative stress **(Fugure 1G)**. We also found that ROS accumulation in BRCA2-depleted cells was comparable between *wild-type* and *TP53*-deficient backgrounds, indicating that TP53 does not contribute to the generation of ROS itself (**Figure S1M**). To evaluate the functional contribution of this oxidative surge, cells were treated with the antioxidant α-tocopherol (α-TOC) (**Figure S1N**). α-TOC treatment effectively suppressed ROS accumulation (**Figure S1N**) and significantly alleviated the growth delay observed in BRCA2-depleted cells (**Figure S1O**). These results suggest that ROS generation is one of the primary causes of the growth suppression induced by BRCA2 depletion.

Furthermore, we validated these findings using the pancreatic cancer and ovarian cancer-derived cell lines, CAPAN1 and PEO1, respectively. Compared with cells harboring *wild-type BRCA2* (*pbBRCA2*) (**Figure S1P**), BRCA2-deficient cells exhibited strikingly higher sensitivity to H_2_O_2_ (**Figures S1Q and S1R**). Fluorescence imaging with ROS probes also revealed prominent spontaneous ROS accumulation in CAPAN1 cells, which was completely abolished by BRCA2 complementation or NAC treatment (**Figure S1S**). These findings suggest that BRCA2-deficient cells are subjected to a persistent oxidative burden, which confers hypersensitivity to exogenous oxidative insults such as H_2_O_2_.

### Accumulation of 8-oxoG induced by ROS in *BRCA2*-depleted genomic DNA

We examined the accumulation of 8-oxoguanine (8-oxoG), one of the most prevalent oxidative DNA lesions induced by ROS^34^, in genomic DNA extracted from cells treated with auxin for 24 h. Using slot-blot analysis, we found that 8-oxoG accumulated in the genomic DNA of auxin-treated *BRCA2*^AID^ TK6 cells, and a similar accumulation was observed in BRCA1-depleted TK6 cells (**Figure 2A**). Since NRF2 is a master transcriptional regulator that coordinates the cellular antioxidant response to counteract oxidative stress, we investigated its role under the BRCA2-depleted condition. Notably, 8-oxoG levels were further elevated in cells depleted of both NRF2 and BRCA2 (**Figure 2A**), indicating that the ROS generated upon BRCA2 loss is normally mitigated by NRF2-dependent antioxidant mechanisms, and that NRF2 contributes to the protection of cells from oxidative DNA damage under conditions of BRCA2 deficiency.

**Figure 2.**
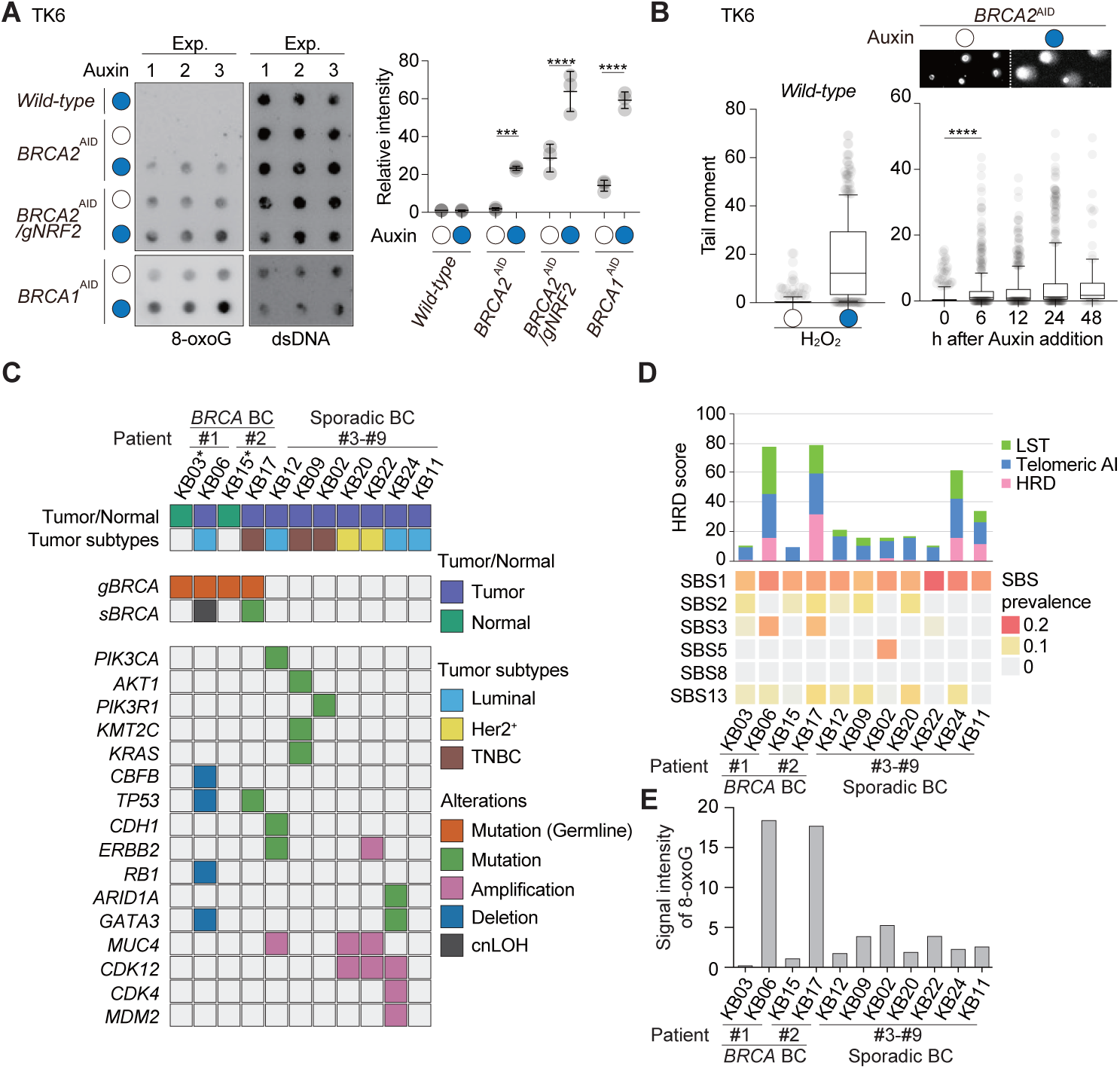
BRCA2 deficiency leads to 8-oxoG accumulation and PARP-dependent repair signaling. (A) Dot blot quantification of genomic 8-oxoG, showing increased oxidative DNA damage in *BRCA1/2*^AID^ and *NRF2*-deficient cells with 500 µM auxin for 24 h. (B) Alkaline comet assay measuring single-strand DNA breaks. Tail moment increases over a 0 – 48 h time course after the addition of 500 µM auxin. (C) The detected pathogenic genetic alterations in *BRCA*-deficient breast tumors and sporadic breast tumors. KB03 and KB15 indicate the contralateral normal mammary glands relative to the tumors in the prophylactically resected specimens. (D) Upper panel: HRD scores of normal and tumor samples for each patient, calculated as the sum of Loss of Heterozygosity (LOH), Telomeric Allelic Imbalance (TAI), and Large-scale State Transitions (LST). Lower panel: scores of the corresponding breast cancer associated SBS signatures for each sample. (E) Quantification of 8-oxoG signal intensities in the clinical samples shown in Figure S3C. Statistical significance was calculated by two-way ANOVA followed by Sidak’s multiple comparisons test (A) or Mann-Whitney test (B). \**p* < 0.05, \*\**p* < 0.01, \*\*\**p* < 0.001, \*\*\*\**p*<0.0001.

### Accumulation of single-strand breaks (SSBs) after BRCA2 depletion

OGG1 excises oxidized bases to generate abasic (AP) sites, which are incised by endonucleases such as APE1, producing single-strand-break (SSB) intermediates. To assess oxidative DNA damage after BRCA2 depletion, we performed alkaline comet assays following auxin-induced degradation in TK6 cells. H₂O₂ (100 µM) treatment produced prominent DNA tails, which validated assay sensitivity (**Figure 2B**). BRCA2-depleted cells exhibited increased tail moments, with similar trends in *BRCA1*^AID^ and *PALB2*^AID^ cells (**Figures 2B, S2A, and S2B**), indicating that loss of BRCA1, BRCA2, or PALB2 elevates oxidative DNA damage. Notably, these oxidative lesions were detectable as early as 6 h after depletion. These findings indicate that BRCA2 depletion triggers a burst of ROS within the first 6 h, establishing a pro-oxidative environment that precedes the manifestation of overt genomic instability. Consistent with an oxidative-damage response, p53 stabilization was detected at 12 h after auxin addition (**Figure S2C**).

### 8-oxoG accumulation in BRCA-deficient clinical specimens

To evaluate the association between BRCA depletion and oxidative DNA damage under physiological condition, 8-oxoG levels in human breast cancer (BC) samples were analyzed. Tumor samples from BC patients, including two hereditary breast and ovarian cancer (HBOC) cases (Patient #1 and #2) carrying *BRCA1/2* mutation and seven sporadic cases, were included in this study. Additionally, for the HBOC cases, non-tumor mammary gland samples were obtained from prophylactic mastectomy specimens **(Figure S3A and S3B**). The detected pathogenic genetic alterations are illustrated in **Figure 2C** and details are provided in **Table S4**. Among the HBOC cases, biallelic *BRCA1* mutations (comprising a germline mutation and copy neutral loss of heterozygosity) were identified in Patient #1. Similarly, biallelic *BRCA2* mutations (comprising germline and somatic mutations) were detected in Patient #2. These findings were consistent with their clinical testing results (Patient #1, *BRCA1* p.Leu63Ter carrier; Patient #2, *BRCA2* p.Tyr2154Ter carrier). Among the sporadic cases, pathogenic mutations in driver genes frequently observed in BC (*e.g*., *PIK3CA*, *AKT*, and *PIK3R1*) or *ERBB2* amplification were identified. High homologous recombination deficiency (HRD) scores were observed in *BRCA-*deficient tumors, but not in non-tumor samples. Meanwhile, only two of the sporadic cases (Patient #8 and 9) showed a similar tendency, indicating non-*BRCA* HRD (**Figure 2D**). Consistent with the established association between HRD and SBS3, SBS analysis showed a prominent SBS3 contribution in *BRCA*-deficient tumors^35–37^. While ROS-derived mutations are physically present, the limits of deconvolution likely preclude the isolation of a discrete ROS-associated signature like SBS18. Instead, these mutational footprints are likely subsumed into SBS3, which may function as an integrated genomic feature reflecting the concurrent processes of DNA repair failure and oxidative stress.

Slot-blot analysis revealed that 8-oxoG levels were markedly elevated in *BRCA-*deficient tumor compared to their respective normal mammary counterparts, suggesting the high 8-oxoG levels were driven by elevated ROS production triggered by the biallelic *BRCA* loss (**Figures 2E and S3C**). The magnitude of oxidative damage in seven sporadic BC cases (Patient #3-10) did not reach the high 8-oxoG signals observed in *BRCA* tumors although there was a trend toward higher 8-oxoG levels compared to the non-tumor mammary gland (**Figure 2E**). These data indicate that 8-oxoG, primarily induced by oxidative stress, is highly and specifically elevated in *BRCA*-deficient tumor compared to *BRCA*-proficient tumor. This suggests that the failure to maintain redox homeostasis may be a distinctive hallmark of *BRCA*-deficient tumors, potentially driving the chromosomal instability observed in these malignancies. Although these clinical observations are limited by a small sample size, the accumulation of 8-oxoG specifically in regions with biallelic *BRCA1*/*2* loss aligns with our findings in controlled cell culture models and confirms its occurrence in clinical settings.

### Quantitative genomic landscape of 8-oxoG accumulation

To define the high-resolution genomic distribution of 8-oxoG and investigate how BRCA2 deficiency reshapes the landscape of oxidative DNA damage, ChIP-seq was performed using a specific antibody against 8-oxoG in TK6 cells (**Figure S4A**). To ensure precise quantification, lambda DNA, which was subject to spontaneous oxidation in aqueous solution, was employed as an internal spike-in control (**Figures S4A and S4B**). This quantitative analysis revealed that while basal 8-oxoG peaks present in control cells due to endogenous oxidative damage, BRCA2 depletion reorganized this genomic landscape.

We identified 72,408 peaks (**Figures S4C and S4D**) common to DMSO and auxin, along with 29,628 DMSO-specific peaks (**Figures S4C and S4E**) and 37,537 auxin-specific *de novo* peaks (**Figures S4C and S4F**) only in *BRCA2*-depleted cells. Consistent with nucleotide preferences, these peaks were predominantly localized within GC rich regions (**Figures S4D-S4F**). Systematic annotation identified a subset of 5,090 genes associated with auxin-specific *de novo* 8-oxoG peaks within their coding regions or were located in the immediate vicinity of these peaks (**Figure S4G**).

To determine the functional relevance of these localized lesions, the relationship between auxin-dependent 8-oxoG accumulation and transcriptomic alterations was evaluated (**Figure S4H**). Comparison of these 5,090 genes with all other genes revealed that those associated with *de novo* 8-oxoG signals exhibited a statistically significant increase in transcript levels following BRCA2 depletion (**Figure S4I**). Collectively, these findings demonstrate that the reorganized landscape of genomic oxidative damage is linked to transcriptional alterations in the absence of BRCA2. Using global expression changes as a background, Pearson’s chi square test identified a significant enrichment of genes with moderate transcriptional induction (log_2_(fold change), 0 to 2) among those harboring *de novo* 8-oxoG peaks (**Figure S4J**). These findings suggest that oxidative DNA damage preferentially accumulates at genomic regions where transcription coupled open chromatin facilitates the access of ROS to DNA.

### Constitutive PARP activation under BRCA loss

Under BRCA2 deficiency, ROS accumulate and oxidative DNA damage persists (**Figures 1 and 2**). We next asked whether DNA repair is constitutively engaged in response to this ongoing damage. We focused on PARP signaling, which is activated during base excision repair. PARP1 and PARP2 bind to SSBs and utilize NAD⁺ as a substrate for auto-poly(ADP-ribosyl)ation. We therefore examined whether PARylation increases when BRCA1 or BRCA2 is depleted in TK6 cells (**Figures 3A and S5A**). Immunoblotting showed elevated PARylation upon BRCA1 or BRCA2 depletion (**Figure 3A**). Inhibition of PARG, which removes poly(ADP-ribose), consequently enhanced the PAR signal (**Figure 3A**). Consistent with these findings, flow cytometric analysis in *BRCA2*^AID^ cells revealed a BRCA2-depletion-dependent increase in PAR signal (**Figure 3B**). These results indicate that, in BRCA-loss cells, ROS are continuously generated and oxidative damage persists, and that PARylation is constitutively engaged to support repair.

**Figure 3.**
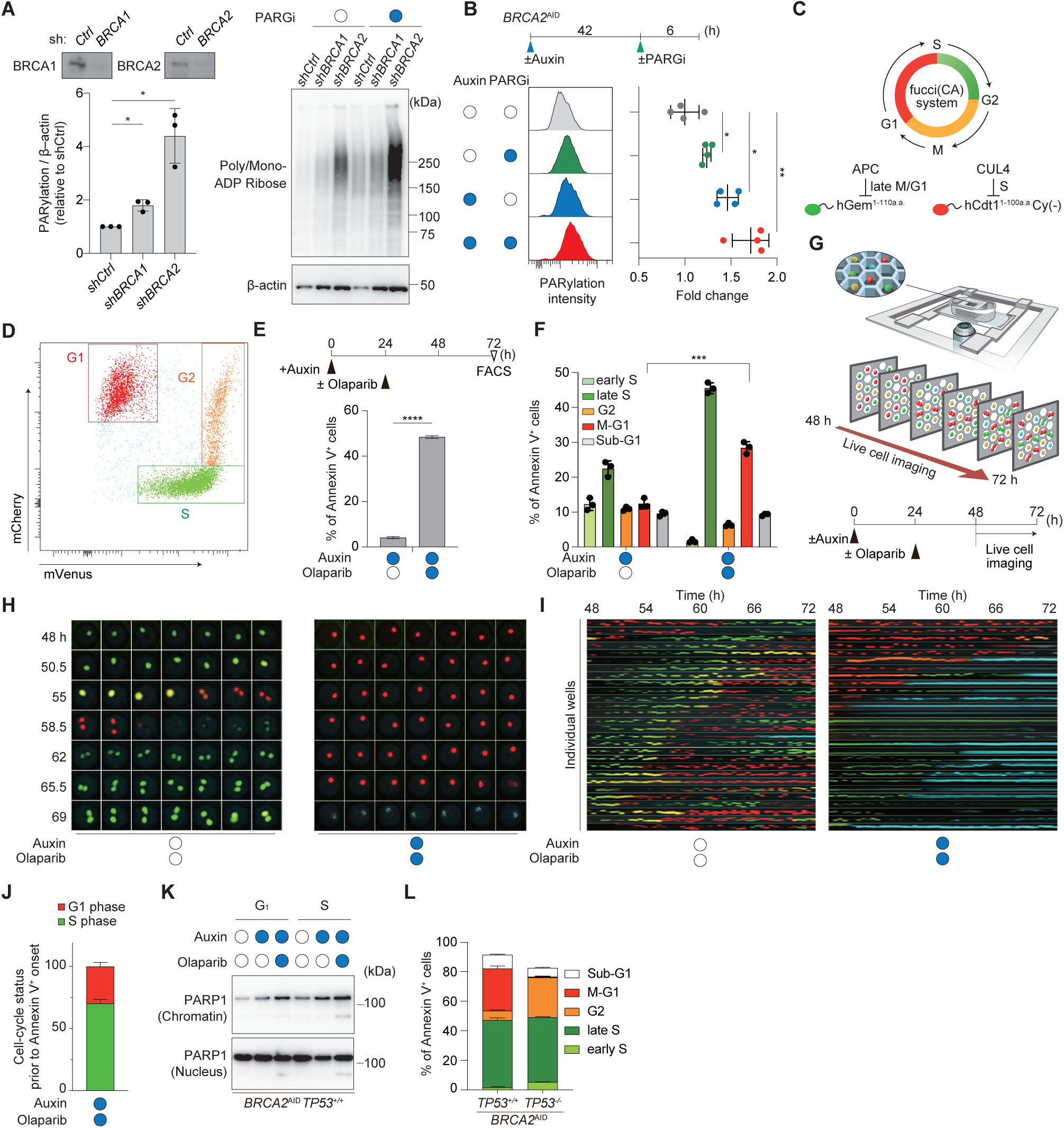
Olaparib-induced PARP1 trapping in G_1_ triggers apoptosis in BRCA2-deficient Cells. (A and B) Mono/poly(ADP-ribose) (PAR) levels in *BRCA1/2*-depleted cells. Immunoblots of sh*RNA*-treated cells exposed to 10 µM PARG inhibitor (PARGi) for 12h (A). FACS analysis for *BRCA2*^AID^ cells treated with 500 µM auxin (48 h) and 10 µM PARGi (6 h) (B). (C and D) Fucci reporter system: schematic (C) and representative FACS plots distinguishing G_1_ (mCherry^+^ and mVenus^−^), S (mCherry^−^ and mVenus^+^), and G_2_/M (mCherry^+^ and mVenus^+^) phases (D). (E) Annexin V⁺ fraction at 72 h after 500 µM auxin ± 10 µM olaparib. (F) Cell-cycle distribution within Annexin V^+^ population by Fucci. (G–I) Live-cell imaging: scheme (G), representative time-lapse images (H), and single-cell tracking images (I). (J) Cell-cycle status immediately before apoptosis onset (∼30% undergo apoptosis directly from G_1_). (K) Chromatin fractionation showing PARP1 trapped on chromatin in sorted G_1_ and S phase cells. (L) *TP53* requirement: Annexin V⁺ distribution in *TP53^+/+^* and *TP53*^−/−^ backgrounds after 500 µM auxin (72 h) + 10µM olaparib (48 h). Data are presented as mean ± SD from at least three-independent experiments. Statistical significance was calculated using an unpaired t-test (A, B, E, and F). \**p* < 0.05, \*\**p* < 0.01, \*\*\**p* < 0.001, \*\*\*\**p* < 0.0001.

### Cell cycle phase-dependent cytotoxicity of olaparib in BRCA2-deficient cells

PARP-mediated repair is chronically elevated under BRCA2 loss, potentially creating vulnerability to PARP inhibitors. We therefore asked whether pharmacologic PARP inhibition exploits this redox-repair liability in a phase-dependent manner across the cell cycle. We therefore quantified olaparib cytotoxicity and mapped the timing of cell death. To delineate the cell-cycle dynamics underlying this vulnerability, we utilized Fucci (CA) reporter system into *BRCA2*^AID^ TK6 cells^38^ (Fucci-*BRCA2*^AID^ cells) (**Figures 3C and 3D**). Following 24 h of auxin-induced BRCA2 degradation, cells were treated with olaparib and analyzed via flow cytometer. In the presence of auxin alone, apoptotic (Annexin V-positive) cells were maintained at low levels (**Figures 3E, left panel in S5B, and S5C**) and distributed across all cell-cycle phases **(Figures 3F and left panel in S5D**). Conversely, the addition of olaparib significantly increased apoptotic fraction (**Figures 3E, right panel in S5B, and S5C**), predominantly in late S phase (**Figures 3F and right panel in S5D**). Notably, we also observed a substantial increase in apoptotic cells arrested in G_1_ phase (**Figures 3F and right panel in S5D**).

To capture the temporal dynamics of the apoptotic process, we performed single-cell live imaging (**Figure 3G**). In control cells, G_1_-phase cells (mCherry-positive; red) progressed through S phase (mVenus-positive; green) and successfully underwent cytokinesis (**left panels in Figures 3H, 3I, S5E, and Video S1**). In contrast, BRCA2-depleted cells treated with olaparib frequently exhibited prolonged G_1_ arrest and underwent apoptosis without entering S phase (**right panels in Figures 3H, 3I, S5E, and Video S2**). Notably, approximately 30% of the cells that became Annexin V-positive had never exited G_1_ phase (**Figure 3J**), confirming that a significant fraction of olaparib-induced cell death originates directly from G_1_.

To elucidate the mechanism driving this G_1_-phase apoptosis, we examined PARP1 trapping on chromatin via Western blot analysis. Using the Fucci reporter system, G_1_-phase cells were isolated with nearly 100% purity, excluding S-phase contamination. We found that PARP1 was significantly enriched in the chromatin fraction following olaparib treatment in both G_1_ and S phases (**Figure 3K**). These suggest that PARP1-DNA complexes formed during G_1_ contribute to an apoptotic response before the onset of DNA replication.

We next investigated the requirement for TP53 in G_1_-specific apoptosis. In *TP53*-deficient (*TP53^−/−^*) cells, the G_1_-phase apoptotic fraction was almost entirely abolished, while the proportion of apoptosis in late S phase and G_2_ phase significantly increased (**Figures 3L and S5F**). These data indicate that the G_1_-phase arrest and subsequent apoptosis induced by BRCA2 depletion and PARP inhibition are strictly dependent on the TP53 pathway.

### Generation and validation of a *Brca2*^AID^ mouse model

To investigate the effect of BRCA2 deficiency *in vivo*, we engineered a murine model incorporating an improved AID system^39^. A mAID tag was inserted in-frame immediately upstream of the endogenous termination codon of the *Brca2* locus, enabling *in vivo* BRCA2 degradation upon administration of the synthetic auxin analog 5-Ph-IAA (**Figures S6A-S6C**). The OsTIR1 (F74G) mutant protein, which confers high-affinity binding to mAID in the presence of 5-Ph-IAA, was expressed ubiquitously under the control of the CAG promoter and targeted to the *Rosa26* locus^40^ (**Figure S6A**). Mice harboring both the homozygous *Brca2*^AID/AID^ alleles and the *Rosa26*^CAG-TIR(F74G)^ transgene were viable and born at expected Mendelian ratios (**Figure S6D**), indicating that the genetic modifications do not perturb embryonic development or early postnatal viability.

### Functional loss of HR and hypersensitivity to DNA crosslinking agents *in vivo*

Intraperitoneal (IP) administration of 5-Ph-IAA induced rapid BRCA2^AID^ degradation within four hours, which persisted for at least 24 hours (**Figure 4A**). To assess HR activity, mice were irradiated following three days of daily 5-Ph-IAA injections, and thymocytes were analyzed for RAD51 foci formation four hours later. RAD51 foci were markedly reduced in 5-Ph-IAA-treated mice compared with controls, indicating that BRCA2 depletion abolishes HR *in vivo* (**Figure S6E**). We next evaluated the impact of BRCA2 loss on cisplatin sensitivity, as repair of cisplatin-induced DNA crosslinks requires HR (**Figure 4B**). Following 5-Ph-IAA administration, we analyzed two genotypes: *Brca2*^AID/AID^ *Rosa26*^+^ (control) and *Brca2*^AID/AID^ *Rosa26*^CAG-TIR(F74G)^ (**Figure S6B**). Low dose cisplatin treatment in control mice did not result in weight loss, depletion of LSK (Lin^−^, Sca-1^+^, c-Kit^+^) cells including hematopoietic stem cells (HSCs) or mortality (**Figures 4B-4D, and S6F**). In contrast, *BRCA2*-depleted mice exhibited severe weight loss, marked reduction of LSK cells, and succumbed within six days of cisplatin exposure (**Figures 4C, 4D, and S6F**). Our data indicate that 5-Ph-IAA induces targeted degradation of BRCA2, thereby abolishing the HR pathway essential for repairing cisplatin-induced DNA crosslinks, establishing functional BRCA2 loss *in vivo*.

**Figure 4.**
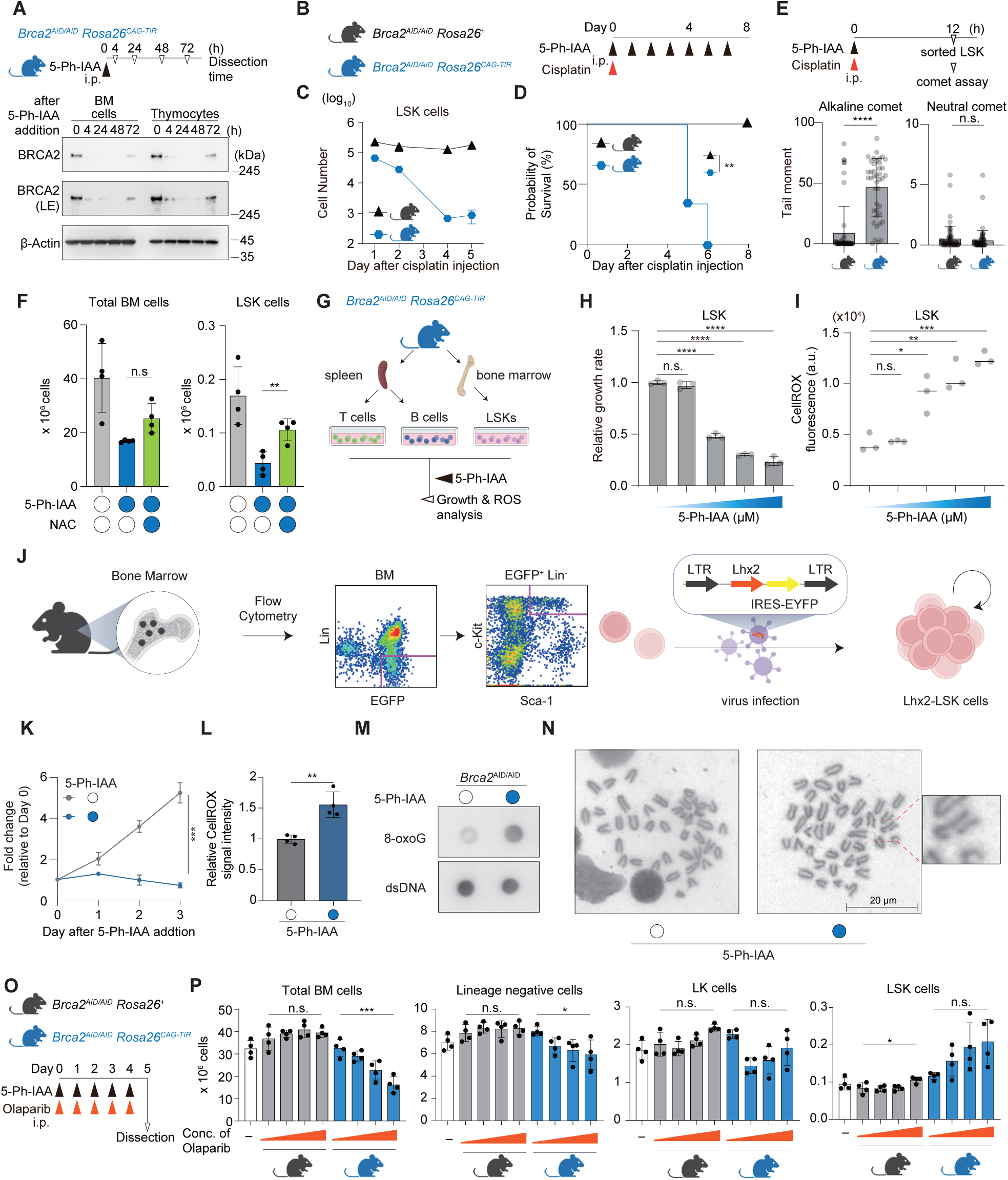
BRCA2 is essential for hematopoietic stem cell maintenance and redox homeostasis *in vivo*. (A) *In vivo* BRCA2 degradation in bone marrow (BM) and thymus. Immunoblot of BRCA2 in BM and thymocytes from the mice at the indicated time points after 5-Ph-IAA administration. (B–D) Hypersensitivity to cisplatin upon BRCA2 depletion: scheme (B), LSK quantification (C), and Kaplan-Meier survival analysis (D) (E) Accumulation of DNA breaks in LSK cells: experimental scheme (upper), alkaline (left) and neutral (right) comet assays in LSK cells isolated 5-Ph-IAA-injected mice (lower). (F) Antioxidant NAC rescues BM and LSK cell numbers in BRCA2-depleted mice. (G–I) *Ex vivo* dose-response to 5-Ph-IAA (0, 0.001, 0.01, 0.1, 1 µM): scheme (G), growth rates (H), and intracellular ROS (I). (J–N) Genomic instability and oxidative damage in Lhx2-LSK progenitors: schematic of Lhx2-LSK generation (J), growth retardation (K), ROS levels (L), 8-oxoG accumulation (M), and metaphase spreads showing chromosomal aberrations (N) after treatment with 100 nM 5-Ph-IAA. Scale bar, 20 µm. (O and P) *In vivo* hypersensitivity to PARP inhibition: experimental timeline (O), quantification of BM subpopulations (P) in *Brca2*^AID^ mice after administration of 5-Ph-IAA and Olaparib. Data are presented as mean ± SD from at least three-independent experiments. Statistical significance was calculated using an unpaired t-test (E, F, H, I, K, L, and P) and Kaplan-Meier survival analysis (D). \**p* < 0.05, \*\**p* < 0.01, \*\*\**p* < 0.001, \*\*\*\**p* < 0.0001.

### Mechanisms of hematopoietic failure: ROS-induced DNA damage in *BRCA2*^AID^ mouse

Having established that BRCA2 depletion *in vivo* leads to a severe defect in HR activity, we next investigated the mechanistic underpinnings of the severe hematopoietic failure observed in our model. Specifically, we sought to determine whether the ROS-driven oxidative stress and single-strand break (SSB) accumulation observed in our mouse model were recapitulated within the hematopoietic stem and progenitor cell (HSPC) compartment *in vivo*. To this end, we performed comet assays on LSK cells isolated 12 hours after the co-administration of 5-Ph-IAA and cisplatin (**Figure 4E**). While neutral comet assays showed minimal changes in tail moments (**right panel in Figure 4E**), indicating that double-strand breaks were not the primary lesion at this early time point, alkaline comet assays revealed a pronounced increase in tail moments (**left panel in Figure 4E**), consistent with a substantial accumulation of SSBs induced by BRCA2 loss (**Figure 2B**). Importantly, administration of N-acetylcysteine (NAC) restored the number of LSK cells (**Figures 4F and S6G**). These findings demonstrate that BRCA2 depletion in LSK cells promotes oxidative stress and subsequent DNA damage.

### Lineage-specific vulnerabilities and genomic instability in hematopoietic progenitors

To further characterize lineage-specific effects, we isolated T cells, B cells, and LSK from *Brca2*^AID/AID^ *Rosa26*^CAG-TIR(F74G)^ mice and cultured them *ex vivo* for four days following 5-Ph-IAA addition (**Figure 4G**). LSK and B cells, but not T cells, exhibited a dose-dependent increase in intracellular ROS upon BRCA2 depletion, accompanied by profound proliferation arrest (**Figures 4H, 4I, and S6H-S6K**). Together, these results indicate that BRCA2 loss in hematopoietic stem and progenitor cells induces oxidative stress, leading to accumulation of single-strand DNA lesions, and that this phenotype can be mitigated by antioxidant treatment.

To examine the impact of BRCA2 depletion on LSK cells, we established Lhx2-induced LSK (Lhx2-LSK) cells by retroviral transduction of Lhx2, a LIM-homeobox transcription factor (**Figure 4J**). The expanded Lhx2-LSK cells retained the Lin⁻Sca-1⁺c-Kit⁺ phenotype (**Left panel in Figure S6L**) and maintained their multilineage myeloid differentiation potential under standard culture conditions^41,42^ (**Right panel in Figure S6L**). 5-Ph-IAA-mediated degradation of BRCA2 markedly abolished the formation of ionizing radiation-induced RAD51 foci formation, thereby validating the functional loss of HR (**Figures S6M and S6N**). Acute BRCA2 depletion profoundly impaired Lhx2-LSK proliferation and led to a marked increase in intracellular ROS (**Figures 4K and 4L**). BRCA2 loss also resulted in substantial accumulation of 8-oxoG (**Figure 4M**) and a significant increase in chromosomal aberrations (**Figure 4N**). Together, these findings demonstrate that BRCA2 depletion in LSK cells not only disrupts HR but also initiates a cascade of metabolic and genomic stress, characterized by impaired proliferation, elevated ROS, accumulation of oxidative DNA lesions, and severe chromosomal instability.

### Effect of BRCA2 depletion on PARP inhibitor-induced myelosuppression *in vivo*

Clinically, hematological toxicities such as anemia and neutropenia are frequent complications of PARP inhibitor therapy and often limit their clinical utility^43–45^. To investigate the mechanisms underlying these toxicities and to assess the impact of olaparib, a PARP inhibitor, on hematopoiesis under BRCA2 deficiency *in vivo*, we daily administrated mice with olaparib and 5-Ph-IAA from Day 0 to Day 4, followed by analysis of bone marrow and peripheral blood on Day 5 (**Figure 4O**). We found that olaparib treatment induced a dose-dependent reduction in total bone marrow cellularity and the lineage-negative cell population in BRCA2-depleted mice (**Figure 4P**). Consistent with this myelosuppression, peripheral blood analysis revealed a progressive decline in white blood cell (WBC) and red blood cell (RBC) counts, as well as hemoglobin (HGB) levels (**Figure S6O**). Strikingly, however, the LSK fraction, enriched for the most primitive hematopoietic stem cells, exhibited a clear dose-dependent expansion in response to escalating olaparib concentrations, in sharp contrast to the overall depletion of differentiated bone-marrow populations (**Figure 4P**). This enrichment possibly reflects a stress-adaptive mechanism triggered by ROS-induced DNA damage or consequent replication stress, wherein the stem and progenitor compartments expand to compensate for impaired lineage output. The LK (Lin^−^, Sca-1^−^, c-Kit^+^) population, representing committed myeloid progenitors one step differentiated from LSK cells, displayed a similar but less pronounced trend (**Figure 4P**).

### BRCA2 depletion enhances mitochondrial oxidative phosphorylation and activity

To investigate the molecular mechanisms of the cellular response to ROS production and oxidative damage following BRCA2 loss, we performed RNA-seq using *BRCA2*^AID^ TK6 cells. Since BRCA2 protein levels drop below detectable limits within 6 hours of auxin treatment (**Figure 1B**) and significant ROS accumulation is observed by 24 hours (**Figure 1G**), we reasoned that this sustained oxidative environment would manifest as broad transcriptional changes at 24 h after auxin addition. To capture these downstream effects at the transcriptional level, we focused our analysis on the 24-hour time point. Gene Ontology analysis revealed a significant enrichment of genes associated with mitochondrial oxidative phosphorylation (**red circle in Figure 5A**).

**Figure 5.**
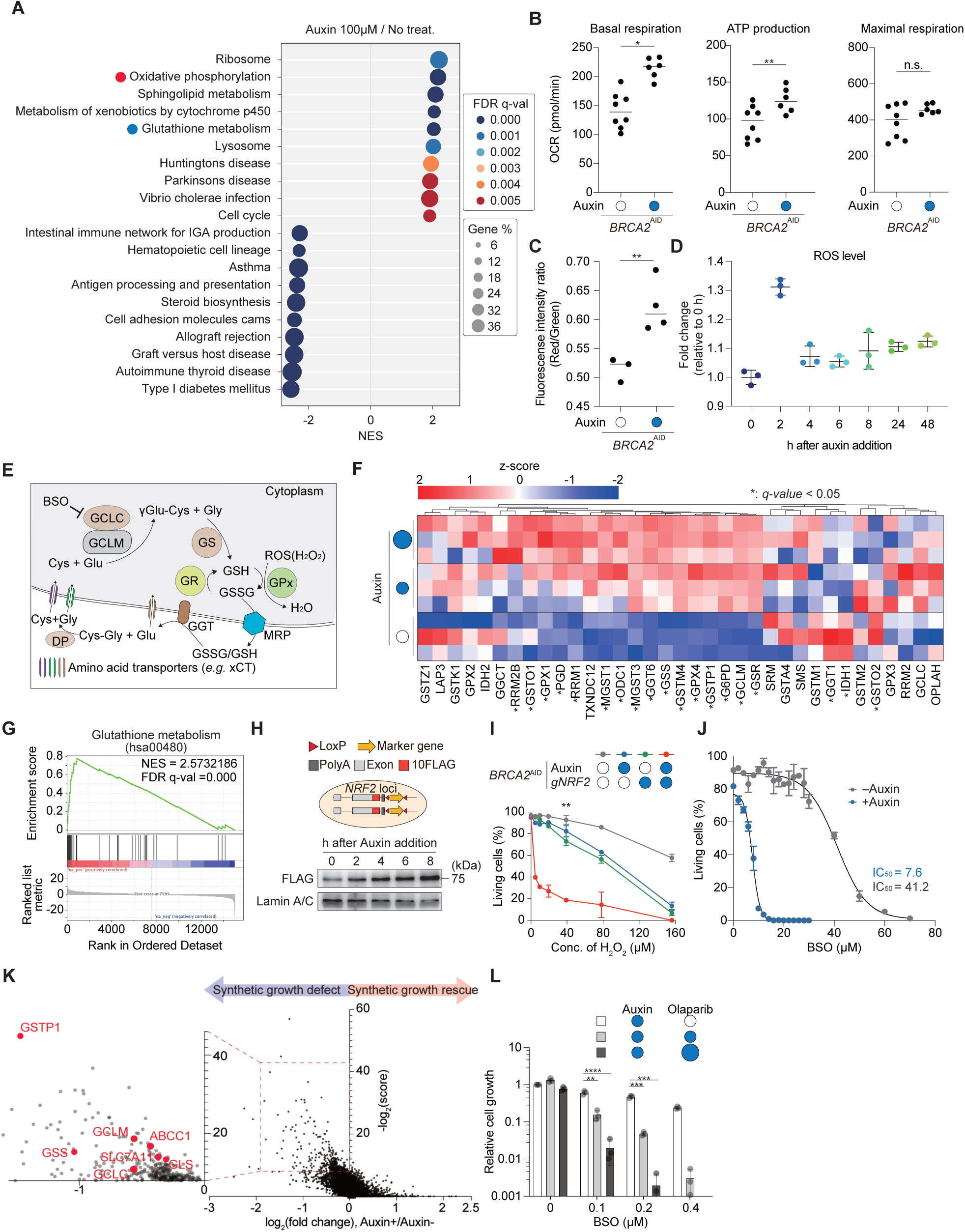
BRCA2 deficiency creates a glutathione-dependent redox vulnerability. (A) Gene Set Enrichment Analysis (GSEA) showing upregulation of glutathione metabolism and oxidative phosphorylation pathways in BRCA2-depleted TK6 cells (100 µM auxin, 24 h) (B) Oxygen consumption rate (OCR) measured using flux analyzer in *BRCA2*^AID^ cells ±500 µM auxin (6 h). (C) Mitochondrial membrane potential (ΔΨm) in cells ± 500 µM auxin (6 h). (D) Intracellular ROS levels following short-term BRCA2 depletion induced by 500 µM auxin. (E) Schematic of glutathione biosynthesis and metabolism. (F and G) Heatmap of gene-expression changes for transcripts annotated to the KEGG glutathione metabolism pathway (hsa00480, KEGG) in cells treated with auxin for 24 h at two doses (100 µM and 500 µM) relative to 0 µM. Values are row z-scores. Asterisks indicate statistically significant differential expression (F). GSEA enrichment plots for hsa00480 comparing 500 µM v.s. 0 µM (24 h). Normalized enrichment score (NES) and FDR *q*-value are indicated for each contrast (G). (H) 10FLAG was inserted into C-terminus of NRF2 locus (upper). Immunoblot showing nuclear NRF2-10FLAG levels and Lamin A/C following short-term BRCA2 depletion induced by 500 µM auxin (bottom). (I) NRF2 depletion sensitizes BRCA2-deficient cells to H_2_O_2_ (dose-response; ±100 µM auxin). (J) BSO hypersensitivity under BRCA2 depletion (±100 µM auxin, 48 h); IC_50_ values indicated. (K) CRISPR screen highlighting glutathione-related genes among synthetic-lethal hits. (L) Synergistic growth inhibition by olaparib and BSO in *BRCA2*^AID^ cells (± 100 µM auxin; olaparib, 0, 2.5, and 5 nM). Data are presented as mean ± SD from at least three-independent experiments. Statistical significance was calculated using an unpaired t-test (B–D, I, and L). \**p* < 0.05, \*\**p* < 0.01, \*\*\**p* < 0.001, \*\*\*\**p*<0.0001

To validate these findings at the functional level, we assessed mitochondrial activity using extracellular flux analysis. We observed increased OCR and ATP production, confirming enhanced mitochondrial respiration (**Figures 5B and S7A**). To mechanistically link this respiratory increase to ROS generation, the mitochondrial membrane potential (ΔΨm) was assessed using the JC-1 at 6 hours post-induction (**Figure 5C**). This early time point was specifically selected to capture the primary mitochondrial alterations immediately following BRCA2 loss. BRCA2-depleted cells exhibited markedly increased JC-1 signal intensity, indicating relative hyperpolarization.

Notably, kinetic measurements of intracellular ROS levels following BRCA2 depletion revealed a biphasic accumulation pattern (**Figure 5D**). We observed a rapid surge in ROS concentration within the first 2 hours of auxin addition, followed by a decline to a plateau level by 4 hours. These findings suggest that the antioxidant defense machinery is activated as a compensatory response to the transient ROS burst. This antioxidant response appears to play a critical role in suppressing the initial spike in ROS levels immediately following BRCA2 depletion.

### Induction of glutathione metabolism and NRF2 protein stabilization following BRCA2 loss

RNA-seq analysis identified glutathione metabolism as another prominent enriched pathway alongside mitochondrial oxidative phosphorylation (**blue circle in Figures 5A, 5E, and 5F**). Gene Set Enrichment Analysis (GSEA) confirmed a significant upregulation of the glutathione metabolism gene set in BRCA2-depleted cells (**Figure 5G**). Western blot analysis further validated these findings: GGT1 was strongly induced following auxin treatment (**Figures S7B**), and GGT1 upregulation was also observed in *BRCA1*^AID^ and *PALB2*^AID^ cells (**Figures S7C and S7D**). Additionally, KGA/GAC, GSS, and xCT were markedly upregulated within 24 h (**Figure S7E**).

In parallel with these metabolic alterations, we observed a marked induction of NRF2, the master transcriptional regulator of the antioxidant response, at the protein level (**Figure 5H**). The accumulation of ROS following BRCA2 loss likely contributes to its stabilization^46,47^, potentially through dissociation from KEAP1. To evaluate the functional significance of this antioxidant response, we generated cells with simultaneous deficiency of NRF2 and BRCA2 and found that their survival rate under oxidative stress, H_2_O_2_, was dramatically reduced (**Figure 5I**).

### Glutathione-dependent redox vulnerability and its therapeutic potential in BRCA2-deficient cells

Our findings indicate that *BRCA2*-deficient cells develop a critical dependency on glutathione to manage acute oxidative stress. Pharmacological assessment using L-buthionine sulfoximine (BSO), a specific inhibitor of the rate-limiting enzyme GCLC in glutathione synthesis **(Figure 5E)**, revealed that BRCA2-depleted cells exhibited higher sensitivity and undergo significant cell death compared to auxin-untreated cells (**Figure 5J**). Similar hypersensitivity was observed in BRCA1– and PALB2-depleted cells (**Figures S7F and S7G**).

Consistent with this observation, our genome-wide CRISPR screen revealed a significant enrichment of genes involved in glutathione synthesis among the synthetic lethal hits required for the survival of BRCA2-deficient cells (**Figure 5K**). Finally, to explore the potential for a therapeutic approach targeting this redox vulnerability, we evaluated the effect of combining olaparib with BSO. The combination treatment killed BRCA2-depleted cells far more effectively than either agent alone (**Figure 5L**). These results suggest that inhibiting glutathione synthesis is a promising strategy to enhance the therapeutic efficacy of PARP inhibitors in *BRCA2*-deficient cancers.

### Establishment of a *Gclm* hypomorphic murine model and early metabolic responses to BRCA2 loss

To investigate the early impact of BRCA2 deficiency on the hematopoietic system, we performed RNA-seq analysis on hematopoietic stem and progenitor cells (LSK cells) 12 hours after acute BRCA2 depletion (**Figure 6A**). GSEA revealed that pathways associated with mitochondrial oxidative phosphorylation and electron transport chain were already significantly enriched at this early time point, reflecting an immediate metabolic response to BRCA2 loss (**Red dots in Figure 6B**). Consistent with this result, 8-oxoG was statistically increased after BRCA2 depletion in Lhx2-LSK cells (**Figure 6C**).

**Figure 6.**
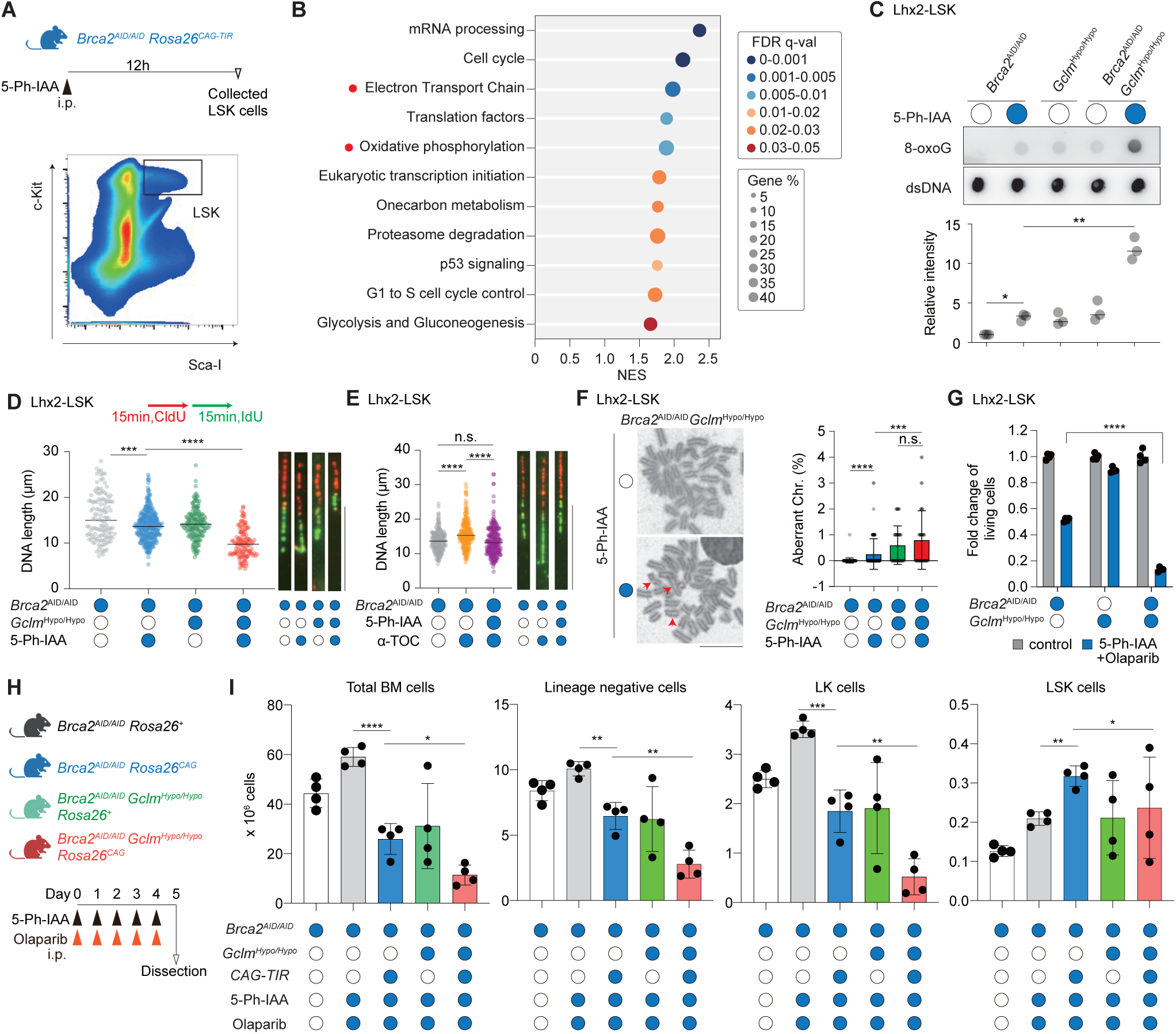
Combined BRCA2 and GCLM deficiency synergistically drives genomic instability and hematopoietic collapse. (A and B) Early transcriptional response 12 h after 5-Ph-IAA: gating of LSK cells (A), GSEA bubble plot using subsets of mouse wikipathways from RNA-seq of LSK cells collected 12 h after 5-Ph-IAA administration. Bubble size reflects gene set size; color encodes FDR *q*-value; position corresponds to the normalized enrichment score (NES) (B). Oxidative phosphorylation and electron transport chain gene set pathways are highlighted with red dots. (C) 8-oxoG dot blot in Lhx2-LSK cells after 1 µM 5-Ph-IAA (6 h); double deficiency (*Brca2*^AID^/*Gclm*^Hypo^ + auxin) synergistically increases 8-oxoG. (D) DNA fiber assay in Lhx2–LSK cells sequentially pulse-labeled with CldU and IdU. Dot plots of nascent DNA tract lengths in BRCA2-depleted, *Gclm*^Hypo^, and BRCA2-depleted/*Gclm*^Hypo^ double-mutant cells (representative fibers are shown). Scale bar, 10µm. (E) Effect of α-tocopherol (α-TOC) on tract length in the indicated genotypes. (F) Chromosomal aberrations in *Brca2*^AID^ *Gclm*^Hypo^ cells treated with 100 nM 5-Ph-IAA (16 h). Scale bar, 10 µm. Red arrows indicate chromosome breaks. (G) Proliferation of Lhx2-LSK cells under partial BRCA2 depletion ± GCLM impairment with olaparib (two days). 4 nM 5-Ph-IAA and 10 nM olaparib were used for BRCA2 depletion. (H and I) Experimental timeline (H) and *in vivo* impact of combined BRCA2 depletion and *Gclm*^Hypo^ on hematopoietic populations after administration of 5-Ph-IAA and olaparib (I). Data are presented as mean ± SD from at least three-independent experiments. Statistical significance was calculated using an unpaired t-test (C–E, G, and I) or Mann-Whitney test (F).

To interrogate the role of glutathione *in vivo* under BRCA2 deficiency, we generated mice carrying a C-terminally AID-tagged *Gclm* allele (*Gclm*^AID^; **Figure S8A**). Even without 5-Ph-IAA, total glutathione was markedly reduced in the liver and erythrocytes (**Figures S8B and S8C**). Despite proper GCLM expression (**Figure S8D**), the presence of the tag appeared to compromise function, causing these mice to phenocopy *Gclm*^−/−^ mice^48,49^ with weight loss (**Figure S8E**). Furthermore, these mice exhibited hypersensitivity to oxidative stress-induced hemolysis following phenylhydrazine (PHZ) administration^50,51^ (**Figures S8F and S8G**). Consequently, we designated this allele as *Gclm*^Hypo^.

### Synergistic accumulation of oxidative damage and replication stress in BRCA2/GCLM-deficient progenitors

To investigate the synergistic effects of BRCA2 depletion and glutathione deficiency, we examined oxidative DNA damage in Lhx2-induced LSK (Lhx2-LSK) cells. Dot blot analysis for 8-oxoG demonstrated that the simultaneous depletion of BRCA2 and impairment of GCLM (*Gclm*^Hypo^) resulted in a striking and synergistic accumulation of lesions that far exceeded the effects of either single perturbation (**Figure 6C**).

To determine if these lesions obstruct replication, we performed single-molecule DNA fiber assays using Lhx2-LSK cells (**Figures 6D and 6E**). While BRCA2 depletion or *Gclm*^Hypo^ alone significantly shortened nascent DNA tract lengths, their combination exacerbated this defect (**Figure 6D**). This shortening was effectively suppressed by the antioxidant α-tocopherol (α-TOC) to the levels observed in the presence of BRCA2 (**Figure 6E**), indicating that ROS act as a primary driver of fork stalling when antioxidant defenses are limited. Crucially, this persistent replication stress was associated with genomic instability. Double mutants displayed an increased frequency of aberrant chromosomes compared to *Gclm*^Hypo^ mutants, a trend that may be confounded by the selective loss of severely damaged cells via apoptosis before metaphase entry (**Figure 6F**). Furthermore, GCLM impairment dramatically potentiated the growth-inhibitory effects of olaparib under BRCA2 depletion. (**Figure 6G**). Collectively, these findings demonstrate that glutathione-mediated antioxidant defense serves as a critical safeguard to prevent ROS-induced replication stress from escalating into catastrophic chromosomal instability and cell death in the absence of BRCA2.

### Synergistic impact of GCLM impairment and BRCA2 depletion on hematopoiesis

To determine whether this synergistic effect is recapitulated in the hematopoietic system *in vivo*, we treated mice with olaparib together with 5-Ph-IAA from day 0 to day 4 (**Figure 6H**). Similar to mice with BRCA2 depletion alone, *Gclm*^Hypo^ mice exhibited myelosuppression following olaparib treatment (**Figure 6I**). This indicates that the detoxification of DNA-damaging agents, including constitutively produced ROS, is essential for maintaining the bone marrow microenvironment and/or cell proliferation. Furthermore, in double-mutant mice harboring both BRCA2 depletion and *Gclm* impairment, we observed severe myelosuppression that far exceeded that of either single condition (**Figures 6I and S8H**), indicating important roles of glutathione in maintenance of bone marrow cells. A striking difference from BRCA2 depletion alone was observed in the LK population (**Figures 6I and 4P**): while this compartment was relatively preserved in single mutants, it underwent a dramatic reduction in the double mutants upon olaparib treatment. Taken together, our findings indicate that glutathione plays a critical role in detoxifying ROS generated by BRCA2 depletion in bone marrow cells, and its deficiency leads to the accumulation of oxidative base lesions. We propose that PARP inhibitors induce cell death independent of cell cycle stages by impeding the repair of these lesions via PARP trapping, thereby blocking the progression of DNA replication and transcription.

### Metabolic reprogramming toward a pro-oxidant state upon BRCA2 loss

Our study demonstrates that BRCA2 deficiency triggers a robust induction of glutathione-related detoxification genes as an adaptive response to the initial surge in ROS. However, ROS-induced DNA damage emerges as early as 6 hours post-depletion and continues to accumulate over time. Additionally, as shown in **Figure 5D**, ROS levels exhibit a progressive increase beginning 24 hours after BRCA2 depletion. To investigate why cells fail to suppress ROS generation and subsequent oxidative genomic damage despite this antioxidant response, we performed comprehensive metabolite profiling in *BRCA2*^AID^ TK6 cells 24 hours after auxin administration **(Table S5).**

Principal component analysis and hierarchical clustering cleanly separated treated from untreated samples, indicating a rapid metabolic shift within 24 h (**Figures S9A and S9B**). Comparative analyses (log₂ transformation and z-score normalization; **see STAR Methods**) revealed a coordinated reduction of antioxidant metabolites (**blue circles in Figures 7A and 7B**) with a reciprocal increase in ROS-promoting species (**pink circles in Figures 7A and 7B**). Together, these data indicate that BRCA2 loss swiftly reprograms metabolism toward a pro-oxidant state, effectively overwhelming the induction of detoxifying genes.

**Figure 7.**
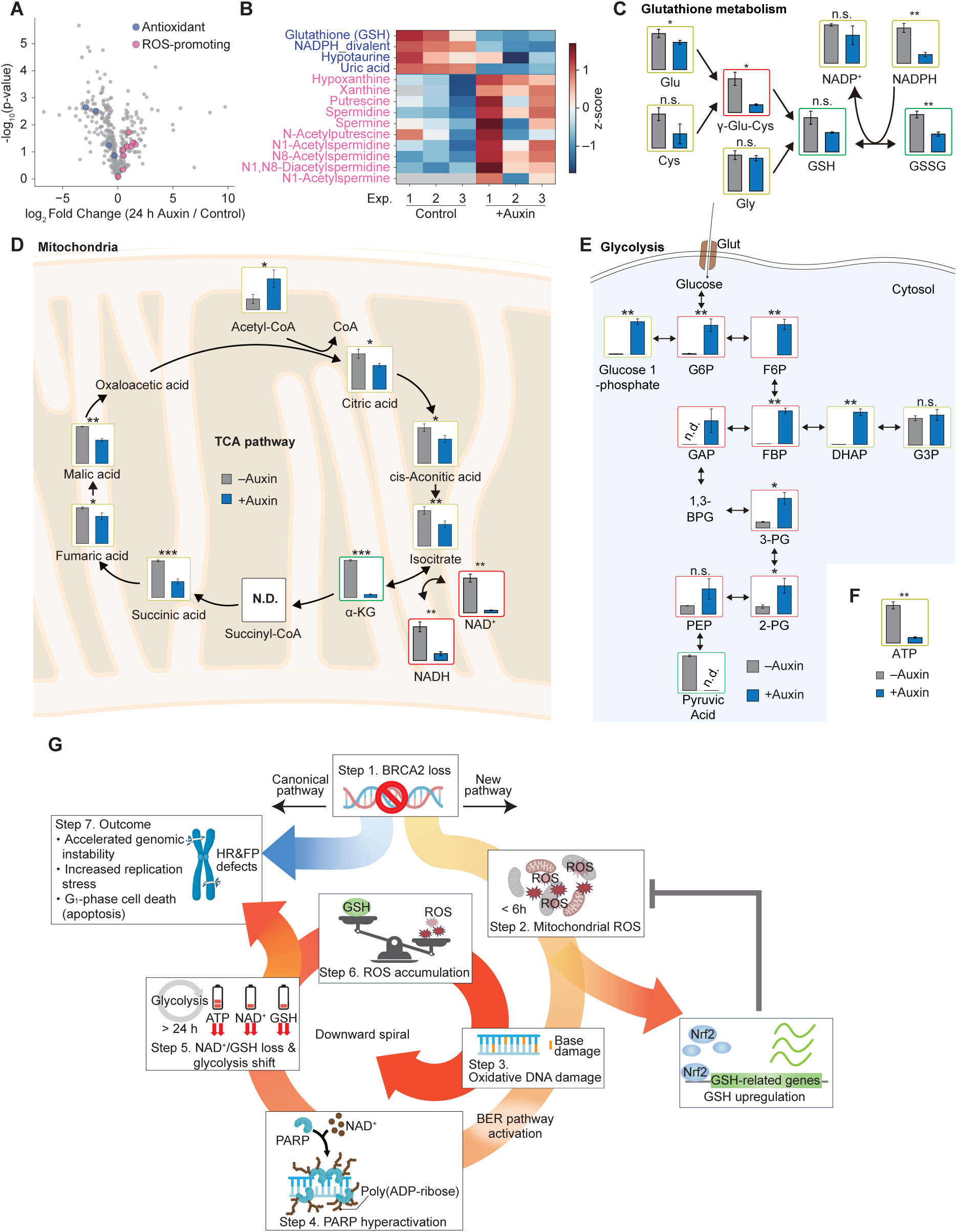
Metabolic rewiring toward a pro-oxidant state and systemic exhaustion of redox cofactors upon BRCA2 loss in TK6 cells. (A) Volcano plot of metabolite changes (auxin/control): pink, ROS-promoting; blue, antioxidant metabolites (B) Heatmap of metabolites in antioxidant/ROS-promoting sets. Colors indicate row z-score. Gray indicates metabolites not detected by mass spectrometry. (C-F) Targeted metabolomics in *BRCA2*^AID^ cells ±auxin: glutathione pathway (γ-glutamylcysteine, GSH/GSSG, cysteine) (C), TCA cycle intermediates (including α-ketoglutarate) and NAD⁺/NADH pools (D), glycolysis metabolites (E), and ATP levels (F). (G) Proposed model summarizing the consequence of BRCA2 depletion. In step 1, the rapid degradation of BRCA2 occurs. As a canonical pathway, this BRCA2 loss immediately induces defects in homologous recombination (HR) and replication fork protection (FP). Rapid degradation of BRCA2 also induces mitochondrial membrane hyperpolarization within 6 h, resulting in increased mitochondrial ROS production (step 2). The elevated ROS simultaneously causes oxidative damage to genomic DNA (step 3) and induces transcription of gene involved in glutathione (GSH) biosynthesis as a compensatory antioxidant response. Single-strand breaks recruit PARP1, leading to excessive PARylation and consumption of NAD^+^ (step 4). NAD^+^ depletion disrupt the TCA cycle, resulting in reduced levels of α-ketoglutarate (α-KG), a key metabolite required for GSH synthesis (step 5). Cells shift toward glycolysis. However, ATP production remains reduced (step 5). Reduced GSH levels impair ROS detoxification, leading to further oxidative DNA damage and establishing a self-reinforcing cycle of oxidative stress (step 6). Ultimately, this cascade results in increased genomic instability, elevated replication stress, and apoptosis in the G1 phase, culminating in cell death (step 7). Data are presented as mean ± SD from at least three-independent experiments. Statistical significance was calculated using an unpaired t-test (C–E). \**p* < 0.05, \*\**p* < 0.01, \*\*\**p* < 0.001, \*\*\*\**p* < 0.0001.

### Systemic exhaustion of glutathione precursors and the NAD^+^ redox pool

Within the glutathione pathway, γ-glutamylcysteine (γ-Glu-Cys) was markedly reduced in BRCA2-depleted cells (**outlined in red, Figure 7C**), accompanied by concomitant decreases in reduced (GSH) and oxidized (GSSG) glutathione (**outlined in green in Figure 7C**). We also detected a pronounced depletion of NAD^+^/NADH pool (**outlined by red, Figure 7D**), which is consistent with the constitutive PARylation observed upon BRCA2 depletion (**Figures 3A, 3B, and S5A**). Although the cystine/glutamate antiporter xCT was upregulated (**Figure S7E**), intracellular cysteine, required for GSH synthesis, was decreased (**outlined in red, Figure S9C**). Homocysteine, a key intermediate in the trans-sulfuration pathway (methionine-to-cysteine biosynthesis), was also substantially reduced (**outlined in green, Figure S9C**), collectively indicating a state of cysteine deprivation in BRCA2-depleted cells. Notably, α-ketoglutarate (α-KG), a metabolite pivotal for both the TCA cycle and glutamate production, declined sharply **(outlined in green in Figure 7D)**. This likely reflects impaired NAD^+^-dependent isocitrate dehydrogenase (IDH) flux due to the severe NAD^+^-dependent crisis. These findings suggest that BRCA2 depletion triggers rapid metabolic reprogramming characterized by impaired redox homeostasis and depletion of essential cofactors, which likely facilitates sustained ROS and persistent DNA damage.

### Glycolytic rewiring and nucleotide collapse in BRCA2-deficient cells

Metabolomic profiling revealed a profound metabolic rewiring following auxin-induced BRCA2 depletion. We observed a robust and generalized accumulation of glycolytic intermediates spanning from glucose-6-phosphate (G6P) to phosphoenolpyruvate (PEP) (**outlined in red, Figure 7E**), whereas pyruvate levels were strikingly depleted (**outlined in green, Figure 7E**). This accumulation within the glycolytic pool was accompanied by a significant redirection of carbon flux into the oxidative branch of the pentose phosphate pathway (PPP), as evidenced by the markedly elevated levels of 6-phosphogluconate (6-PG) and ribulose-5-phosphate (Ru5P) (**outlined in red, Figure S9D**).

However, this upstream expansion of carbon pools does not translate into an increase in downstream anabolic products. In pentose phosphate pathway, while the levels of ribose-5-phosphate (R5P) and phosphoribosyl pyrophosphate (PRPP) remain relatively maintained (**outlined in green, Figure S9D**), there is a drastic and specific depletion of the ADP-ribose pool (**outlined in blue, Figure S9D**). This striking depletion of ADP-ribose likely mirrors the profound collapse of the cellular NAD^+^ pool. Furthermore, the global depletion of nucleoside triphosphates (ATP, GTP, etc.) and deoxynucleoside triphosphates (**Figures 7F and S9E**) suggests that the accumulation of glycolytic and early-PPP metabolites represents a compensatory, yet insufficient, metabolic response to a severe cellular energy crisis and nucleotide exhaustion.

Depletion of deoxyribonucleoside triphosphates (dNTPs) severely compromises DNA replication fidelity and progression, leading to replication stress and accumulation of DNA lesions^52–55^. Given that TCA cycle dysfunction limits ATP production, we hypothesized that BRCA2-depleted cells rely heavily on glycolysis for survival. To test this, we cultured cells in galactose-containing medium, forcing cells to rely on mitochondrial oxidative phosphorylation^56^. BRCA2-depleted cells exhibited a markedly more pronounced growth defect relative to untreated controls (**Figure S9F**). These findings indicate that BRCA2 depletion impairs mitochondrial respiratory capacity, forcing a metabolic shift toward glycolysis that fails to compensate for the underlying energy and nucleotide deficit.

Taken together, BRCA2 deficiency persistently drives mitochondrial ROS-initiated oxidative damage and reliance on base excision repair and, through PARP hyperactivation with depletion of NAD^+^ and glutathione and a shift toward glycolysis, establishes a self-reinforcing repair-metabolism loop that underlies olaparib cytotoxicity across cell-cycle phases including G_1_ and hematopoietic toxicity, as demonstrated in cell systems and in mice.

## Discussion

### BRCA2 regulates redox-metabolic networks to prevent a downward spiral of genomic and metabolic collapse

Our study demonstrates that BRCA2 loss instigates a self-reinforcing redox-repair loop that couples mitochondrial physiology to genome maintenance beyond canonical failure of HR and replication fork protection (FP). As illustrated in our model, the loss of BRCA2 initiates a multi-step cascade characterized by a distinct temporal sequence (**Figure 7G**). This spiral begins with the loss of BRCA2, which compromises HR and replication fork protection (Step 1). Within 6 hours, we observed a primary trigger consisting of a surge in mitochondria-derived ROS driven by mitochondrial hyperactivation (Step 2). This intensified oxidative stress leads to the accumulation of 8-oxoG lesions in transcriptionally active chromatin and engages the base excision repair (BER) pathway. Our genome-wide screen reinforces this link by identifying a critical genetic dependency on BER factors for the survival of *BRCA2*-deficient cells, whose absence results in the buildup of hazardous single-strand-break (SSB) intermediates (Step 3). Consequently, PARP1 becomes hyperactivated (Step 4). By 24 hours, the persistent demand for repair precipitates a systemic metabolic crisis characterized by the exhaustion of NAD^+^ and glutathione pools along with a shift toward glycolysis (Step 5). This metabolic state promotes further accumulation of ROS, which feeds back to Step 3 by further amplifying oxidative DNA damage and SSB accumulation (Step 6). Ultimately, the convergence of genomic instability and metabolic exhaustion triggers TP53-dependent apoptosis in the G_1_ phase (Step 7).

### Mitochondrial overactivation as the root cause of *BRCA2*-deficient phenotypes

Our metabolomics reveal why the antioxidant response fails to contain ROS after BRCA2 loss. PARP hyperactivation drives a collapse of the NAD⁺ pool, cysteine and γ-glutamylcysteine fall despite xCT upregulation, and α-ketoglutarate decreases with constrained TCA flux, which together throttle glutathione synthesis and redox recycling. ATP and dNTP pools drop and cells shift toward glycolysis, yet this compensatory rewiring does not restore cofactor or nucleotide availability. As a result, detoxification capacity is uncoupled from demand and ROS generation outpaces repair (**Figure 5D**), explaining the early onset and continued accumulation of oxidative genomic damage despite robust induction of glutathione-related genes.

Importantly, this self-reinforcing spiral was not limited to cell culture models but was faithfully recapitulated in our murine hematopoietic system, where mitochondrial ROS-induced oxidative stress exacerbated the genomic instability inherent to HR deficiency, collectively precipitating severe bone marrow failure. While BRCA2 deficiency manifests through an exceptionally diverse array of complex phenotypes, ranging from hematopoietic stem cell loss and chromosomal instability to G_1_-specific apoptosis, our findings highlight the critical importance of the antioxidant response and demonstrate that these disparate events converge on a complex interplay between canonical HR failure and mitochondrial overactivation, with the latter acting as a potent driver of the resulting surge in ROS production. These results position BRCA2 as a dual-function regulator that integrates DNA repair with the modulation of mitochondrial respiration to prevent the excessive production of organelle-derived ROS^57^. By acting as a key regulator of these redox-metabolic networks, BRCA2 ensures that cellular metabolic activity does not outpace genomic repair capacity, a concept tightly linked to NAD^+^-dependent mitochondrial homeostasis and redox control^58^. Therefore, we propose that the preservation of genomic integrity by BRCA2 is inextricably linked to its novel function in restraining mitochondrial overactivation. This idea provides a unifying mechanistic framework for understanding BRCA-deficient oncology and establishes a compelling rationale for therapeutic strategies that pair PARP inhibition with redox modulation.

### A pan-cell-cycle model for PARP-inhibitor cytotoxicity

We challenged the conventional view that the cytotoxicity of PARP inhibitors is restricted to S-phase replication stress^11,12^. In this study, we showed that olaparib induces *TP53*-dependent apoptosis in G_1_ (**Figure 3**). On the basis of our observation, we propose a pan-cell-cycle model. In this model, ROS-induced single-strand breaks (SSBs) and persistent BER intermediates render chromatin throughout the cell cycle intrinsically susceptible to PARP trapping, committing cells to apoptosis. When these lesions are not resolved before entry into S phase and are compounded by a profound NAD⁺ deficit that limits repair capacity and cellular energetics, cytotoxicity is amplified, culminating in TP53-dependent G_1_ arrest and apoptosis. These findings suggest that, in addition to combination therapies targeting S-phase vulnerabilities, such as ATR-CHK1 pathway inhibition and PARP inhibitors, therapeutic strategies targeting redox rewiring represent a promising approach.

### Limitations of the Study

While this study identifies BRCA2 as a critical integrator of redox homeostasis and metabolic rewiring, several limitations warrant consideration. First, our findings rely on an acute auxin-inducible degron (AID) model to capture immediate mitochondrial and ROS responses. Although this approach effectively reveals early triggers, chronic BRCA2 deficiency in clinical settings may involve distinct adaptive mechanisms through long-term metabolic reprogramming or compensatory repair pathways. Furthermore, while we utilized cell line and murine models, tissue-specific metabolic environments vary across human organs, requiring further characterization beyond the hematopoietic system. Second, the precise molecular targets through which BRCA2 regulates mitochondrial respiration remain unidentified. It is unclear whether BRCA2 directly modulates the respiratory chain or if these effects arise indirectly from nuclear signals of genomic integrity. Elucidating this direct interaction is essential for understanding the metabolic liabilities of *BRCA*-deficient tumors. Third, clinical translation of our proposed therapeutic strategies requires careful optimization. Although combining PARP inhibitors with redox modulators effectively targets *BRCA2*-deficient cells, normal tissues like bone marrow also exhibit high sensitivity to oxidative stress. Future research must refine the therapeutic index to maximize tumor lethality while minimizing healthy tissue toxicity. Fine-tuning the timing and intensity of such interventions will be critical for effective clinical implementation.

## Acknowledgements

We thank all members of the Genome Dynamics project and of the Tokyo Metropolitan Institute of Medical Science (TMIMS) for their technical assistance and helpful discussions. This research was supported by JSPS KAKENHI Grant, 19H04267; Takeda bioscience Research grant; Naito Research grant; Astellas research foundation (to HS), Multilayered Stress Diseases (JPMXP1323015483), Science Tokyo, Kose Cosmetology Research grant, and the Sasakawa Scientific Research Grant from The Japan Science Society. This work is also supported by a grant for Special Research from the Tokyo Metropolitan Government since 2025.

## Author Contributions

M.T. and H.S. conceptualized the study. M.Y., H.Sh., M.T.K., and I.T. developed the methodology. Investigation was performed by H.H., T.I., K.K., K.Ya., Ko.T., Y.K., M.S., R.K., Ko.Y., S.Y., A.E., D.S., N.L., M.Y., H.Sh., T.K., Y.N., R.S., N.T.T.T., S.N., J.M., K.H., K.Yu., and H.S. Formal analysis was conducted by H.H., T.I., K.K., K.Ya., Ko.T., Y.K., M.S., R.K., Ko.Y., S.Y., A.E., D.S., Ka.T., N.L., R.S., H.K., and H.S. Resources were provided by S.W., H.M., N.N., T.K., T.T., S.N., M.T.K., I.T., K.H., H.K., and K.Yu.. H.H. and H.S. wrote the original draft; all authors reviewed and edited the manuscript. H.S. supervised the study and managed project administration. M.T. and H.S. acquired the funding.

## Declaration of interest

The authors declare no competing interests.

## STAR METHODS

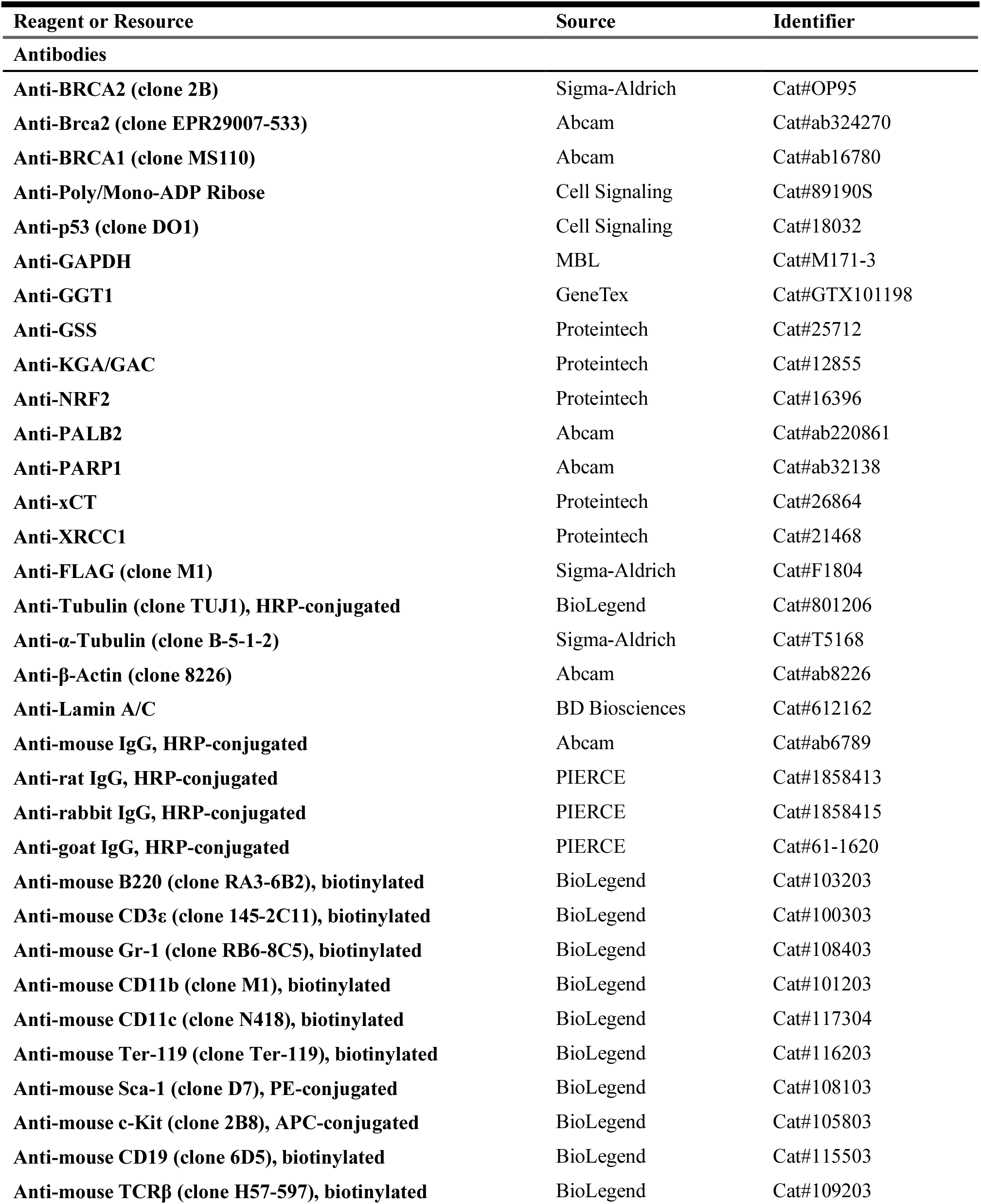
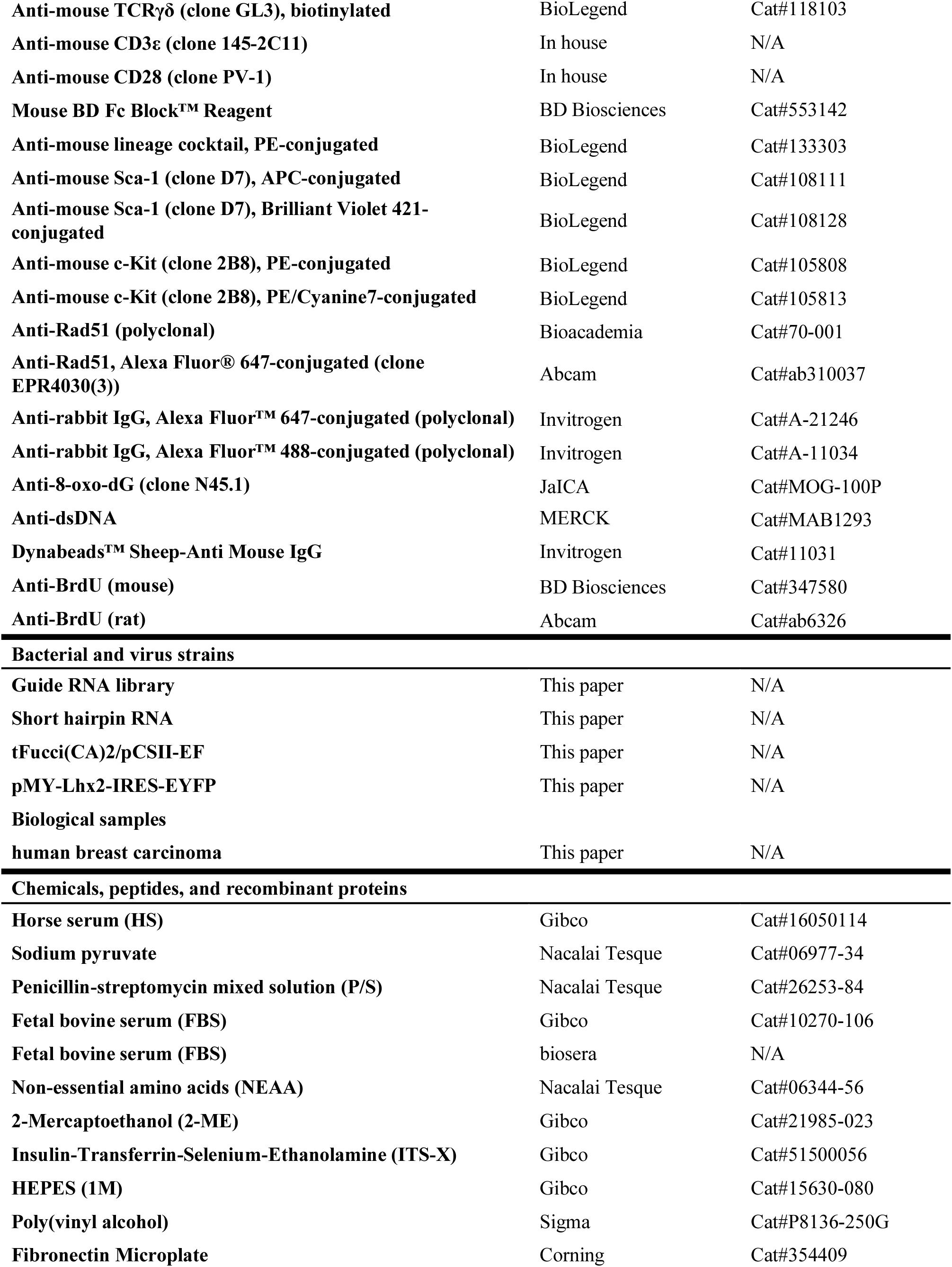
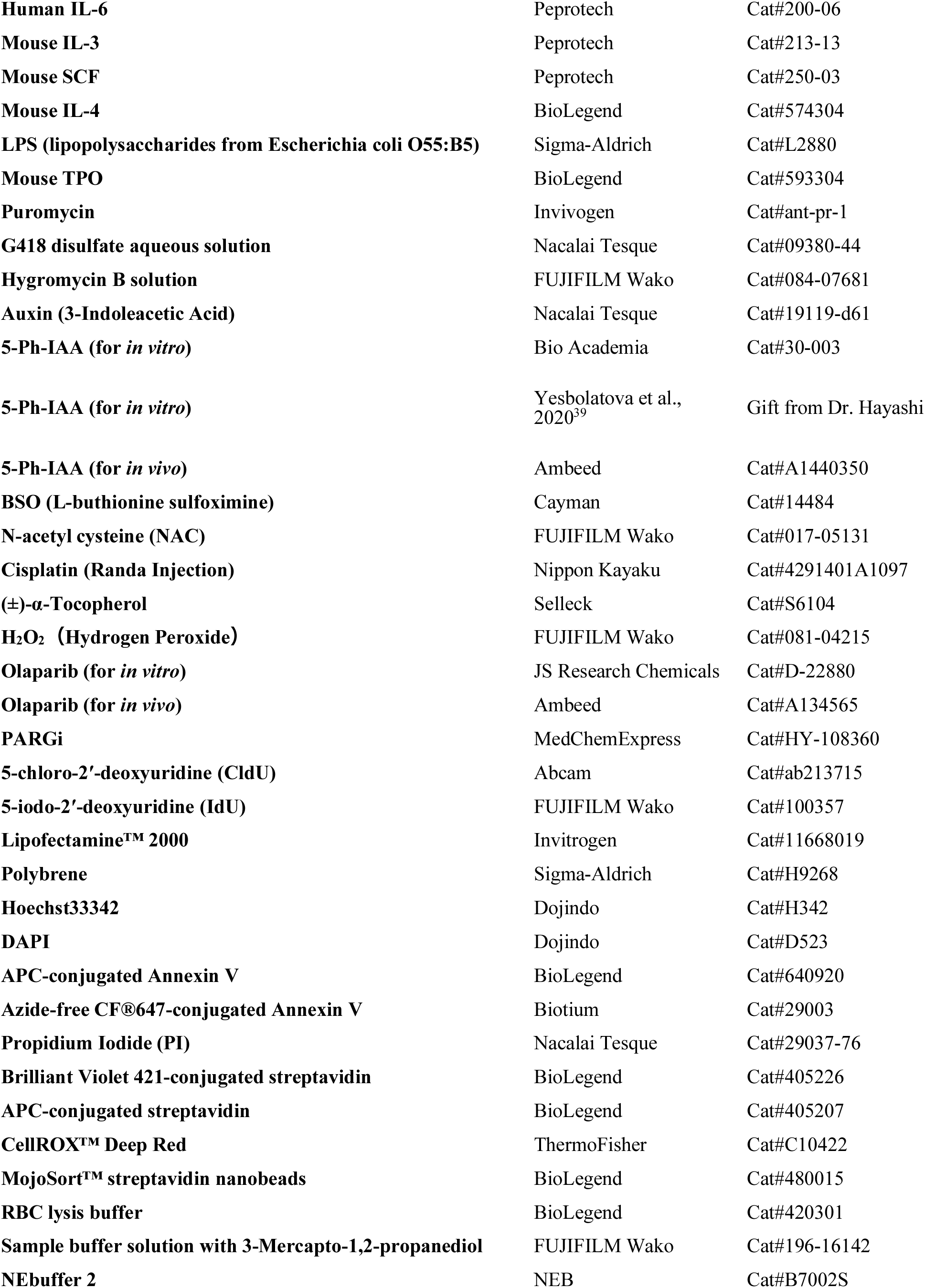
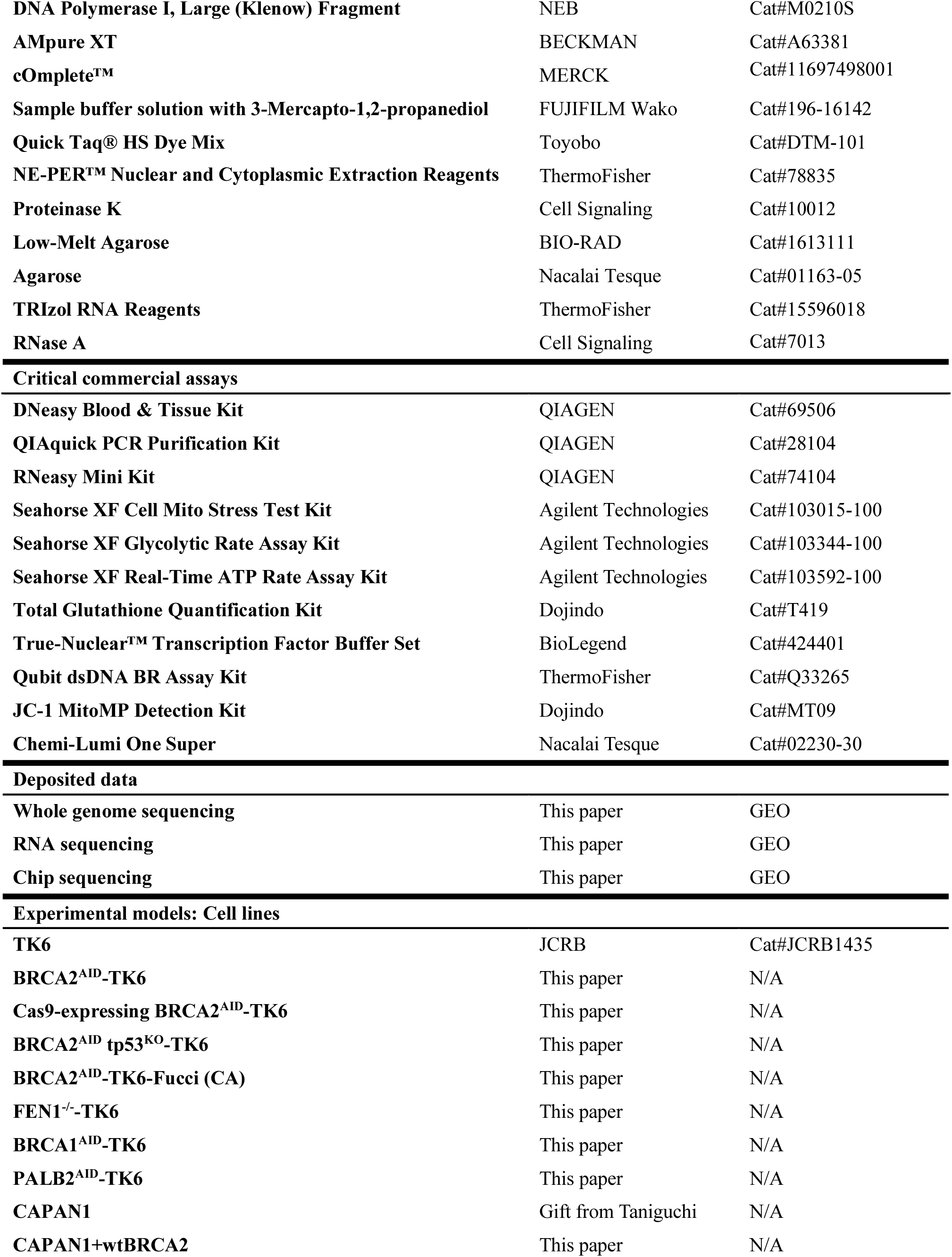
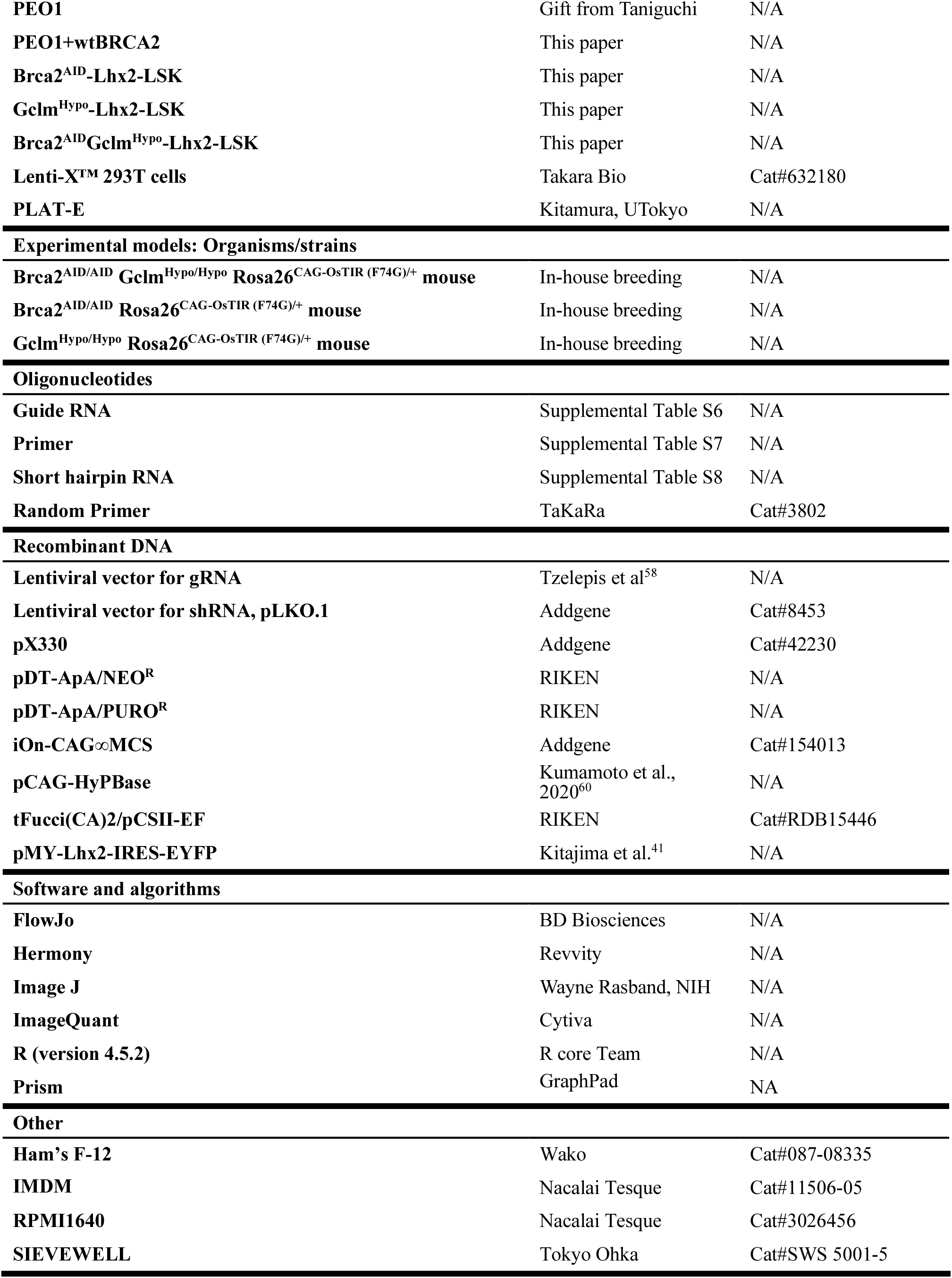
KEY RESOURCES TABLE.

## EXPERIMENTAL MODEL AND STUDY PARTICIPANT DETAILS

### Cell lines and culture conditions

Human B cell line TK6 was obtained from Japanese Collection of Research Bioresources Cell Bank (JCRB1435; Osaka, Japan). TK6 and its derivatives were grown in RPMI1640 medium (3026456, Nacalai Tesque) supplemented with 5% horse serum (HS, 16050114, Gibco), 200 mg/mL sodium pyruvate (06977–34, Nacalai Tesque), and penicillin-streptomycin mixed solution (P/S, 26253-84, Nacalai Tesque). *BRCA2*-deficient human cell lines, CAPAN1 pancreatic adenocarcinoma cell line and PEO1 ovarian cancer cell line were cultured in IMDM (11506-05, Nacalai Tesque) supplemented with 20% fetal bovine serum (FBS, 10270-106, Gibco) and P/S, RPMI1640 supplemented with 10%FBS and P/S, respectively. Lenti-X™ 293T cells and PLAT-E cells were grown in DMEM supplemented with 10% FBS and P/S. Immortalized LSK cells, named as Lhx2-LSK cells, were kept in IMDM supplemented with 10% FBS (biosera), non-essential amino acids (NEAA, 06344-56, Nacalai Tesque), 55 µM 2-Mercaptoethanol (2-ME, 21985-023, Gibco), P/S, 20 ng/mL human recombinant interleukin-6 (IL-6, 200-06, Peprotech), and 100 ng/mL mouse recombinant stem cell factor (SCF, 250-03, Peprotech).

### Human tumor tissues

For prophylactic mastectomy, paired normal and breast cancer tissues were collected from patients during surgical treatment or biopsy. All experiments in this study were approved by both the Research Ethics Committees of the Tokyo Metropolitan Institute of Medical Science (Approval No. 25-16) and Tokyo Metropolitan Komagome Hospital (Approval No. 3211).

### Mice

All animal care and experimental procedures were approved by the Institutional Animal Care and Use Committee of the Tokyo Metropolitan Institute of Medical Science (Approval No. 25−036) and performed in accordance with institutional guidelines. Wild-type C57BL/6J mice were purchased from CLEA Japan, Tokyo, Japan. Mice were housed in a specific pathogen-free (SPF) facility under a standard 12-hour light/12-hour dark cycle at a controlled temperature of 23 ± 1°C and 50 ± 10% humidity. The *Brca2*^AID^ (*Brca2*^AID/AID^*Rosa26*^CAG-OsTIR/+^) mouse and *Gclm*^Hypo^ (*Gclm*^Hypo/Hypo^*Rosa26*^CAG-OsTIR/+^) mouse lines were established in this study (see below).

## METHOD DETAILS

### Genetic manipulations of cell lines

To establish a cell line expressing OsTIR1, TK6 cells were co-transfected with the targeting vector pMK231 carrying with OsTIR1-IRES-EGFP and the CRISPR/Cas9 plasmid pX330 carrying with gRNA for AAVS1 locus. At 48 hours post-transfection, the cells were subjected to selection with puromycin (final concentration: 1 µg/mL) to isolate stable integrants. Genotyping PCR were performed to confirm the targeting event.

*BRCA2*^AID^-TK6 cells were generated by CRISPR/Cas9 system. For CRISPR/Cas9 gene editing, guide RNA was inserted into the expression vector pX330 (42230, Addgene) carrying human codon-optimized SpCas9. The guide RNA sequences were listed in Supplemental Table S6. To construct targeting vectors, the left and right homologous arms were amplified from genomic DNA by PCR. These fragments were inserted into DT-ApA/NEOR or DT-ApA/PUROR vector (RIKEN) and co-transfected with pX300 vector expressing corresponding guide RNAs into TK6-derived cells. The newly established cells were verified by genomic PCR. The list of primer sequences was listed in Supplemental Table S7.

For transduction of *BRCA2*, *BRCA1,* or control short hairpin *RNA* (*shRNA)* into wild type and *FEN1*^−/−^-TK6 cells, the lentiviral vector carrying shRNA was co-transfected with virus packaging plasmids into Lenti-X™ 293T cells by Lipofectamine™ 2000 (11668019, Invitrogen) according to manufacturer’s instruction. Then, supernatants were collected 48 hours after the transfection and transduced into TK6-derived cells. Sequences for shRNA were shown in Supplemental Table S8. The transduction of Fucci (CA2) into *BRCA2*^AID^-TK6 cells was carried out using a lentivirus vector CSII-CMV-MCS-IRES2-Venus. The transduced cells were cloned by limiting dilution and *BRCA2*^AID^-TK6-Fucci (CA2) cells were established.

Enforced expression of *wild-type* (*wt*) human BRCA2 (NCBI Gene ID: 675) into CAPAN1 and PEO1 cells was carried out by Piggy Bac transposon system using iOn-CAG∞MCS (154013, Addgene) and pCAG-HyPBase vectors. These vectors were co-transfected using Invitrogen Neon® Transfection System with the following parameters: pulse voltage, 1500 V; pulse width, 10 ms; pulse number, 3.

### CRISPR/Cas9 screening

This pooled CRISPR/Cas9 knockout screen is meticulously designed to uncover genetic vulnerabilities, specifically synthetic lethal interactions, that emerge following acute loss of BRCA2 function within the Cas9-expressing BRCA2^AID^ –TK6 cell line. Initially, cells are transduced with a comprehensive lentiviral gRNA library at a low MOI, ensuring single gRNA integration per cell, and robust library coverage is strictly maintained throughout. A five-day selection with Puromycin is subsequently performed to establish a stably gRNA-expressing cohort. The selected population is then split into two arms: a Vehicle Control Group and a Selection Group treated for five days with 40 µM Auxin (19119-61, Nacalai Tesque). Under this auxin concentration, BRCA2-AID protein levels are undetectable by Western blotting analysis (**Fig. S1C**), and the overall cellular proliferation rate remains comparable to that of *wt* cells (**Fig. S1D**). This Auxin treatment thus generates a state of specific BRCA2 deficiency without imposing a non-specific growth arrest, acting as a selective pressure that causes the preferential depletion of cells harboring gRNAs that target genes synthetically lethal with the isolated BRCA2 loss. Following selection, genomic DNA is isolated from both groups, and the gRNA cassettes are amplified via nested PCR before being quantified using deep Next Generation Sequencing (NGS). Finally, bioinformatic analysis calculates the log-fold change (LFC) in gRNA representation between the Auxin-treated and control groups, definitively nominating the most depleted gRNAs and their corresponding target genes as essential factors for survival under compromised BRCA2 function.

### RNA-seq analysis

RNA extraction from TK6 cells was performed by RNeasy Mini Kit (74104, QIAGEN). Paired-end RNA-seq was performed in BRCA2^AID^-TK6 cells under three conditions, 0 µM, 100 µM, and 500 µM auxin, with three biological replicates for each condition. Raw FASTQ files were aligned to the human reference genome (GRCh37) using HISAT2 with the ––dta option and known splice sites extracted from the corresponding Ensembl GTF annotation (release 87) supplied via the ––known-splicesite-infile option. For mouse primary LSK cells, sequencing reads from two biological replicates for each condition were aligned to the mouse reference genome (GRCm39) using HISAT2 with default parameters. Gene-level read counts were obtained using featureCounts from the Subread package. Gene annotations GRCh37.87.gtf (human) or GRCm39.111.gtf (mouse) were used as reference annotation files. The resulting count matrix was then imported into DESeq2 for differential expression analysis. After DESeq2 normalization and dispersion estimation, log2 fold changes and p values were calculated using the Wald test, and Benjamini–Hochberg adjusted p values (padj) were obtained. Variance-stabilizing transformation (vst) was applied to the count data, and the transformed values were used for heatmap visualization.

For gene set enrichment analysis (GSEA), the Wald test statistic from the DESeq2 results table was used as the ranking metric. GSEA was performed in PreRanked mode using the GSEA desktop software with KEGG gene sets (c2.cp.kegg_legacy.v2025.1.Hs.symbols.gmt and m2.cp.wikipathways.v2026.1.Mm.symbols.gmt), and normalized enrichment scores (NES) and FDR q-values were calculated. Genes belonging to the KEGG glutathione metabolism pathway (KEGG_GLUTATHIONE_METABOLISM) were extracted from the KEGG GMT file, and the corresponding gene symbols were matched to the Ensembl-to-symbol–converted vst expression matrix. For each gene, expression values were standardized across samples by z score (mean = 0, variance = 1), and a heatmap of 0 µM, 100 µM, and 500 µM auxin, arranged in this order, was generated using the pheatmap package. Within this pathway, genes that showed significant differential expression in the comparison between 0 µM and 500 µM auxin (padj < 0.05 in the DESeq2 analysis) were considered significantly regulated and were marked with an asterisk in the heatmap.

### 8-oxoG-IP

Genomic DNA from 5 × 10^6^ cells was extracted with DNeasy Blood & Tissue kit (69504, QIAGEN) following the manufacturer’s instructions. The genomic DNA was sheared to 500 bp average by Branson Digital Sonifier 250D (TIME = 15 sec, AMPLITUDE = 15%, four times, with a three-minute interval on ice). 5 µg genomic DNA in 100 µL was denatured at 100 °C for 15 min and on ice. 100 µL of 4× PBS, 200 µL of 2% BSA, and anti-8-oxo-dG antibody (MOG-100P, JaICA) captured on Dynabeads^TM^ Sheep Anti-Mouse IgG (11031, Invitrogen) and incubated for 16 hours at 4 °C. The beads were washed with Lysis buffer I [50 mM Hepes-KOH pH7.5, 140 mM NaCl, 1 mM EDTA, 0.1% Sodium Deoxycholate, 1% Triton X-100] twice, Lysis buffer II [50 mM Hepes-KOH pH7.5, 140 mM NaCl, 1 mM EDTA, 0.1% Sodium Deoxycholate, 1% Triton X-100], Hight salt Lysis Buffer I [50 mM Hepes-KOH pH7.5, 500 mM NaCl, 1 mM EDTA, 0.1% Sodium Deoxycholate, 1% Triton X-100] twice, Wash Buffer [10 mM Tris-HCl pH8.0, 250 mM LiCl, 1 mM EDTA, 0.5% NP-40, 0.5% Sodium Deoxycholate] twice, and ice-cold TE pH8.0 once. DNA was eluted with 50 µL of Elution Buffer [1% SDS, 50 mM Tris-HCl pH8.0, 10 mM EDTA] from magnetic beads. The immunoprecipitated DNA were treated with 200 µg/mL Proteinase K for 2 hr at 50 °C. The immunoprecipitated DNA were purified by QIAGEN PCR purification kit (28104, QIAGEN) following the manufacturer’s instructions. Immunoprecipitated 8-oxoG DNA and 1 µL of 50 µM Random Primer (3802, TaKaRa) in 44.5 µL volume were denatured at 95 °C for 5min and on ice for 3 min. Denatured DNA were mixed with 5 µL of 10× NEBuffer 2 (B7002S, NEB), 0.5 µL of 10 mM dNTPs, and 1 µL of 5 U/µL Klenow fragment (M0210S, NEB) and incubated at 37 °C for 30 min and inactivated at 75 °C for 30 min. DNA was purified with 45 µL AMPure XP beads (A63381, BECKMAN) and eluted with 50 µL of 0.1× TE.

### Next generation sequencing (NGS)

The input and the immunoprecipitated DNAs were fragmented to an average size of approximately 400 bp by Covaris Focused-ultrasonicator S220. The fragmented DNAs were end-repaired, ligated to sequencing adapters and amplified according to the protocol of NEBNext Ultra II DNA Library Prep Kit for Illumina and NEBNext® Multiplex Oligos for Illumina® (96 Unique Dual Index Primer Pairs, NEB). The amplified DNA was sequenced to paired-end 150 bp reads using illumina sequencer.

### ChIP-seq analysis

For read mapping, a hybrid reference genome consisting of the human genome (GENCODE, GRCh38 primary assembly) and λ DNA (GenBank: J02459.1) was constructed. Sequencing reads were aligned to the hybrid genome using Bowtie2 with the following parameters: –q 30 –F 2308. To perform spike-in normalization, the number of reads mapped to the host genome and λ DNA were calculated for each sample. A spike-in normalization factor (scale factor) was derived from the ratio of λ DNA–mapped reads across samples. Normalized bigWig files were generated using deepTools with the calculated scale factors. Peak calling was performed using MACS2 with the following parameters: –f BAMPE –g hs ––broad ––broad-cutoff 0.01 –q 0.01. Notably, peak calling was performed using IP samples only without input controls. Biological replicates were first assessed for reproducibility by correlation analysis. After confirming high concordance between replicates, peak regions from each replicate were merged using bedtools merge to generate unified peak sets. Correspondingly, merged signal tracks were generated using deepTools bigwigCompare. Based on the merged peak sets from DMSO-treated and auxin-treated samples, peaks were categorized into three groups: (1) peaks detected only in DMSO-treated samples, (2) peaks detected only in auxin-treated samples, and (3) peaks shared between both conditions. To visualize signal enrichment, heatmaps and metaprofiles were generated using deepTools computeMatrix in reference-point mode with the parameters ––binSize 50 and ––missingDataAsZero. For GC content analysis, GC percentages for each peak region were calculated using bedtools nuc with the GRCh38 genome sequence as reference. The calculated GC values were converted into bigWig format for visualization alongside 8-oxoG signal tracks. To identify genes associated with condition-specific 8-oxoG peaks, peak annotations were performed using HOMER (Hypergeometric Optimization of Motif EnRichment) with the annotatePeaks.pl command to obtain gene lists corresponding to peak regions. To assess whether genes harboring 8-oxoG peaks were preferentially distributed across gene expression change upon BRCA2 depletion, genes were classified into four groups based on log2 fold change values (< –2, –2 to 0, 0 to 2, and > 2). The number of genes with or without 8-oxoG peaks in each category was summarized in a contingency table. Independence between gene expression categories and 8-oxoG peak presence was tested using Pearson’s chi-squared test.

To evaluate deviations from expected distributions, standardized Pearson residuals were calculated as:

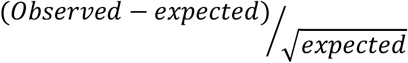

Positive residuals indicate enrichment of genes harboring 8-oxoG peaks relative to expectation, whereas negative residuals indicate depletion.

### Immunoblotting and immunostaining

Whole-cell lysates were prepared from 1×10⁶ TK6-derived cells treated with or without 500 µM auxin for 24 hours or transduced with guide RNAs targeting XRCC1 and NRF2. Cells were lysed in sample buffer containing 3-mercapto-1,2-propanediol (196-16142, FUJIFILM Wako) and sonicated. *FEN1*-deficient TK6 cells transduced with or without shRNAs targeting *BRCA2* and *BRCA1*, as well as CAPAN1 and PEO1 cells with or without enforced expression of wild-type BRCA2, were processed similarly. Protein samples were resolved on 5–20% gradient SDS-Tris-glycine gels and transferred to PVDF membranes. Membranes were probed with primary antibodies listed in the KEY RESOURCES TABLE, including those against *BRCA2*, *BRCA1*, *XRCC1*, *NRF2*, *PALB2*, *PARP1*, *TP53*, and loading controls such as Tubulin, α-Tubulin, GAPDH, and β-Actin. HRP-conjugated corresponding secondary antibodies were used for detection. Chemiluminescent signals were visualized using Chemi-Lumi One Super reagent (02230-30, Nacalai Tesque) and captured with GE Healthcare LAS4000 imaging system using ImageQuant software.

Chromatin fractionation was performed using a commercial chromatin isolation kit, NE-PER™ Nuclear and Cytoplasmic Extraction Reagents (78835, ThermoFisher), according to the manufacturer’s protocol, followed by immunoblotting as described above.

For immunofluorescence staining, cells were collected onto microscope slides using a Shandon Cytospin 4 centrifuge (Shandon), fixed with 4% formaldehyde for 10 minutes at room temperature, permeabilized with 0.5% Triton X-100, and blocked with 0.5% BSA. Alternatively, True-Nuclear™ Transcription Factor Buffer Set (424401, BioLegend) was used for fixation and permeabilization. Cells were stained with antibodies listed in the KEY RESOURCES TABLE. Unconjugated antibodies were visualized using AlexaFluor488– or AlexaFluor647-conjugated secondary antibodies. Nuclear DNA was counterstained with DAPI or Hoechst 33342. Images were acquired using confocal microscopies (Leica SP8/STELLARIS or Revvity Operetta CLS system).

### Detection of poly(ADP-ribose) (PAR) and GGT1 by flow cytometry

Cells were fixed with 4% formaldehyde for 15 min, washed with PBS, and permeabilized with 90% ice-cold methanol for 10 min. After washing again with PBS, cells were incubated with a primary antibody, followed by incubation with an Alexa Fluor 594–conjugated secondary antibody. Fluorescence intensity was measured using an LSRFortessa™ flow cytometer (BD Biosciences), and the resulting fluorescence signal was used as an indicator of intracellular PAR levels. The expression levels of GGT were analyzed by LSRFortessa™ flow cytometry using FITC-conjugated anti-GGT1 antibody (GTX101198, GeneTex).

### Measurement of mitochondrial membrane potential using JC-1

Mitochondrial membrane potential was measured using the JC-1 MitoMP Detection Kit (MT09, Dojindo) according to the manufacturer’s instructions. Cells were suspended in RPMI 1640 medium containing 2 µmol/L JC-1 working solution and incubated for 30 min. After incubation, cells were washed with RPMI 1640 medium and resuspended in imaging buffer solution. Fluorescence signals were analyzed using an LSRFortessa™ flow cytometer, and both green and red JC-1 fluorescence were detected. Mitochondrial membrane potential was quantified as the ratio of red to green fluorescence intensity (Red/Green ratio).

### Cell growth, viability, and stress response assays

Intracellular reactive oxygen species (ROS) levels were measured in CAPAN1 cells with or without enforced BRCA2 expression using 10 µM DCFH-DA and confocal microscopy (Leica SP8/STELLARIS). TK6-derived cells and Lhx2-LSK cells were incubated with 5 µM CellROX™ Deep Red (C10422, ThermoFisher) for 30 minutes and analyzed by LSRFortessa™ flow cytometry.

To detect 8-oxoguanine (8-oxoG) in Lhx2-LSK and TK6, cells were suspended in 1 mL buffer A [10 mM HEPES, pH 7.9; 10 mM KCl; 1.5 mM MgCl₂; 0.34 M sucrose; 10% glycerol; 0.1% Triton X-100; 1 mM PMSF (P7626, SIGMA); protease inhibitor cocktail (11873580001, Roche)] and incubated on ice for 10 min. Ten percent of the suspension (∼100 µL) was reserved for DNA quantification. After centrifugation (1300 × g, 4 °C), the pellet was resuspended in 500 µL buffer B (2 mM EDTA; 0.3 mM EGTA; 0.1% Triton X-100; protease inhibitor) and incubated on ice for 30 min. The suspension was centrifuged (1700 × g, 4 °C, 10 min) to isolate chromatin. The chromatin pellet was washed six times with buffer C (25 mM Tris-Cl, pH 8.0; 800 mM NaCl; 10% glycerol; protease inhibitor) to remove chromatin-associated proteins, then dissolved in 6 M guanidinium chloride. Genomic DNA from human specimens was extracted as described below.

The purified genomic DNA was spotted onto PVDF membranes, blocked, and probed with anti-8-oxoG antibody. Detection was performed using HRP-conjugated secondary antibodies and Chemi-Lumi One L reagent, with imaging via LAS4000 and ImageQuant software.

Alkaline comet assays were conducted to assess DNA damage at the single-cell level. Cells (5×10⁴) were embedded in 1% low-melting agarose and layered onto agarose-coated slides. After lysis (2.5 M NaCl, 100 mM EDTA, 10 mM Tris, 1% Triton X-100), samples were treated with alkaline buffer (300 mM NaOH, 1 mM EDTA), electrophoresed, and stained with ethidium bromide. DNA damage was quantified by measuring tail length and intensity using ImageJ software (https://imagej.nih.gov/ij/docs/faqs.html).

Cell proliferation and survival/viability were assessed by the 3-(4,5-dimethylthiazol-2-yl)-2,5-diphenyltetrazolium bromide (MTT) colorimetric assay, hemocytometer, Coulter counter, or methylcellulose colony formation assay (Yamazaki et al., 2024). Cell viability was also scored using LSRFortessa™ flow cytometry following PI and/or Annexin V staining. The survival of CAPAN1 and PEO1 cells in response to Olaparib treatment was assessed by colony formation assay. Briefly, cells were seeded at low density and treated with indicated concentrations of Olaparib for 18 days.

For DNA fiber assays the cells were sequentially labeled for 15 min with 25 μM 5-chloro-2′-deoxyuridine (CldU, ab213715, Abcam) and 250 μM 5-iodo-2′-deoxyuridine (IdU, 100357, FUJIFILM Wako). The cell concentration was adjusted to 200 cells/µL with PBS and lined 3 µL of cell suspension on the slide and air-dried. The cells were lysed with 60 µL Lysis buffer (0.5% SDS, 200 mM Tris-HCl [pH7.4], 50 mM EDTA) for 5 min. The slides were tilted slightly to drop the lysis buffer down and air-dried. The slides were fixed with fresh Carnoy solution (MeOH:Acetic Acid = 3:1) for 3 min and air-dried. The slides were washed and with –20 °C 70% Ethanol overnight. The slides were washed with PBS 3 times, and denatured in 2.5 M HCl for 30 min at r.t and then were washed with PBS 3 times. The slides were treated with 2 µL of BD™ Purified Mouse Anti-BrdU, clone: B44 (34750, BD Biosciences) and 0.67 µL of Anti-BrdU antibody, clone: [BU1/75 (ICR1)] (ab6326, Abcam) in 100 µL of IMMUNO SHOT immuno-staining, Fine (CSR-IS-F-20, Cosmo Bio) for 1 hr. Then, the slides were washed with PBS-T and treat with 1µL of Alexa488-conjugated anti-mouse antibody (A-21121, ThermoFisher) and 0.25 µL Cy3-conjugated anti-rat antibody (712-165-153, Jackson ImmunoResearch LABORATORIES INC) in 100 µL of IMMNO SHOT immuno-staining, Fine for 1 hr. Then, the slides were washed with PBS-T and mounted with Fluoromount/Plus™ (K048, Diagnostic BioSystems).

The images were obtained using a KEYENCE BZ-X700 microscope equipped with 20 × objective lens. Fiber length was measured by ImageJ. At least 200 bicolor fibers (from CldU to IdU) were counted on each slide. Measurements were recorded from areas of the slides with untangled DNA fibers to prevent the possibility of recording labeled patches from tangled fiber bundles. DNA fiber data (rate of IdU/CldU or IdU length) were represented by dot plot. All statistical comparisons of DNA fiber lengths were performed using the two-tailed, nonparametric, unpaired Mann–Whitney U test, and statistical significance was indicated where applicable.

Oxygen consumption rate (OCR) and extracellular acidification rate (ECAR) were measured using the Seahorse XFe96 Analyzer (Seahorse Bioscience) according to the glycolysis stress test protocol.

### Quantification of total glutathione concentration

*Brca2*^AID/AID^*Gclm*^Hypo/Hypo^*Rosa26*^CAG-OsTIR/+^ and *Brca2*^+/+^*Gclm*^Hypo/Hypo^*Rosa26*^CAG-OsTIR/+^ mice were used as *Gclm* hypomorphic mice. *Wild-type* mice were used as control. Serum, peripheral blood, liver, and bone marrow samples were collected as follows. For serum samples, blood is collected from the mouse heart into BD Microtainer® Blood Collection Tubes (SKU:365967, BD Biosciences) containing a clot activator and a plasma separation agent. Then the tubes were centrifuged at 10,000 rpm for 5 minutes at 24°C. The supernatant was transferred to new tube and 0.25 volume of 10% sulfosalicylic acid in H_2_O (10% SSA) was added. For peripheral blood samples, blood was collected from the mouse heart into MiniCollect® II EDTA-2K tubes (450532, JACLasS). Then 100 µL of blood was transferred to new 1.5 mL tube, centrifuged at 1,000 x g for 10 minutes at 4°C. The supernatant was removed and 200 µL of 10% SSA was added. For liver samples, the liver was excised from the mouse, and 0.15-0.17 g of liver was used for the experiment. 5% SSA was added at a volume of 5 mL per gram of liver and then liver was homogenized. For bone marrow samples, 1 × 10⁶ bone marrow cells were centrifuged at 200 x g for 10 minutes at 4°C. Then the supernatant was removed and the cell pellets were washed with 300 µL PBS (-), centrifuged at 200 x g for 10 minutes at 4°C, and the supernatant was removed. Next, 80 µL of 10 mM HCl was added to the cells and the cells were lysed by two cycles of freezing and thawing. Then, 10 µL of 10% SSA was added.

The samples from serum, peripheral blood, liver and bone marrow were centrifuged at 8,000 x g for 10 min at 4°C and the supernatant was collected. Then, H_2_O was added to adjust the concentration of SSA to 0.5% and used as for the measurements. The measurements were carried out using total glutathione quantification Kit (T419, Dojindo) as manufactures instructions.

### DNA extraction from human tumor specimen

Following surgical resection and subsequent pathological diagnosis to delineate tumor boundaries, tissue specimens were collected using a dermal punch (diameter: 4 mm; length: 8 mm). For normal samples, the mammary glandular components were selectively isolated from the excised breast tissue to ensure high cellular purity. The isolated tumor or glandular fragments were placed in 400 µL of lysis buffer (50mM Tris-HCl pH 7.5, 200mM NaCl, 5 mM EDTA, and 0.2 % SDS) supplemented with Proteinase K (2 µg/mL) and RNase A. The samples were incubated at 56 °C, overnight to ensure complete tissue digestion. For DNA purification, the lysates were subjected to two rounds of phenol/chloroform/isoamyl alcohol extraction, followed by a single extraction with chloroform to remove residual phenol. Genomic DNA was then precipitated by adding sodium acetate and ethanol. The resulting DNA pellets were resuspended in 100 µL of TE buffer. DNA concentrations were determined using the Qubit dsDNA Assay Kit (Q33265, ThermoFisher) on a Qubit Fluorometer according to the manufacturer’s instructions.

### Whole-exome sequencing and variant calling

Genomic DNA was extracted from clinical specimens using the Proteinase K lysis method. Libraries for whole-exome capture were prepared using the Agilent SureSelect Human All Exon V6 kit and subsequently sequenced on an Illumina NovaSeq X Plus system as 150 bp paired-end reads. Mutation discovery was conducted as previously described^61^. Briefly, the obtained reads were trimmed using Trimmomatic to remove low-quality bases and adapter sequences, and then mapped to the human reference genome (hg19) using the BWA-MEM algorithm. Base quality score recalibration and subsequent mutation detection were performed with GATK utilities. Variants were identified using MuTect2 and filtered based on population prevalence to exclude germline background variants. Further filtering was applied with the FilterMutectCalls standard parameter with F-score optimization to ensure sensitivity and specificity. The detected variants were annotated for pathogenicity based on information from the COSMIC and CLINVAR databases. Mutational signatures were identified using the Sigminer R package.

### Copy number alteration analysis and HRD score

CNA analysis was conducted using PureCN against a pool of normal controls. This pool of normals was constructed using data from non-tumor samples lacking specific CNAs, enabling tumor-only CNA analysis. B-allele frequencies were also analyzed using PureCN, which integrates copy number data and variant allele frequence to estimate allelic status, including loss of heterozygosity (LOH) and/or copy-neutral LOH. Tumor purity and ploidy estimated from the CNAs and allelic status were fitted to high-tumor-purity and near-diploid model. Using these data, the homologous recombination deficiency (HRD) score, derived from the number of LOH regions, telomeric allelic imbalance events, and the large-scale state transitions, was calculated using the scarHRD R package.

### Metabolic profiling

*BRCA2^A^*^ID^-TK6 cells were treated with or without 500 µM auxin for 24 hours. Cells were collected, washed, counted, and subjected to methanol extraction. Supernatants were filtered and stored at –80°C. Metabolomic analysis was performed using CE-FTMS (Capillary Electrophoresis–Fourier Transform Mass Spectrometry) in both cation and anion modes by Human Metabolome Technologies (HMT, Yamagata, Japan). A total of 518 metabolite peaks were detected (281 cationic, 237 anionic) across three biological replicates per condition.

Metabolomics data were analyzed using relative abundance values (Relative Area) for each metabolite in Control (n = 3) and Auxin-treated (n = 3) samples. Values below the limit of detection were treated as missing (NA). To avoid undefined values during log transformation, a pseudocount defined as one-half of the smallest positive value observed across the dataset was added to each measurement, and data were log_2_-transformed as log_2_(x + pseudocount). For each metabolite, the log_2_ fold change (log_2_FC) between Auxin-treated and Control groups was calculated from group means as:

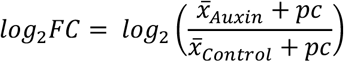

Group differences were assessed using Welch’s t-test (two-sided; unequal variances assumed) on the untransformed relative area values. Metabolites for which the number of valid measurements was < 2 in either group were not assigned a *p* – value (set to missing). Volcano plots were generated with log_2_FC on the x-axis and –log_10_(*p*-value) on the y-axis. Metabolites with missing log_2_FC or *p*-values were excluded from volcano plotting.

Heatmaps were generated for metabolites included in the antioxidant set and ROS-promoting set. For visualization, log_2_-transformed values were standardized on a per-metabolite basis (row z-score), calculated by subtracting the mean across the six samples and dividing by the corresponding standard deviation. Missing values (NA) were retained and displayed in gray.

### Genomic PCR

Genomic DNA was extracted using the DNeasy Blood & Tissue Kit (69506, QIAGEN) and amplified using Quick Taq® HS DyeMix (DTM-101, Toyobo) on a Veriti thermal cycler (Life Technologies). Primer sequences are listed in Supplemental Table S7.

### Single-cell live imaging

*BRCA2*^AID^-TK6-Fucci (CA) cells were treated with 500 µM auxin for 24 hours, followed by co-treatment with 500 µM auxin and 10 µM olaparib for an additional 24 h. Cells were suspended in phenol red-free RPMI-1640 (06261-65, Nacalai Tesque) supplemented with 10% horse serum (16050-122, Gibco) and stained with 0.25 µg/mL CF®647-conjugated Annexin V (29003, Biotium).

Single-cell imaging was performed using SIEVEWELL™ nano-well devices, pre-treated according to the manufacturer’s protocol. Cell suspensions (50,000–70,000 cells) were loaded into SIEVEWELL™ chambers (SWS5001-5, Tokyo Ohka) and imaged using Revvity Operetta CLS system. Time-lapse imaging was conducted every 30 minutes for 24 hours at 37°C in 5% CO₂, capturing nine fields per time point with a 10× objective lens. Data were analyzed using Harmony software.

### Generation of AID-mice

In order to knock-in DNA sequences coding AID tag on *BRCA2* gene (NCBI Gene ID: 12190), double-stranded DNA, that contained 300 bp DNA sequence upstream of the stop codon of *Brca2* gene, the DNA sequence encoding the AID tag, and 300 bp DNA sequence downstream of the stop codon of *Brca2* gene, was synthesized. Then, this DNA fragment was transduced into fertilized oocytes from C57BL/6JJcl (CLEA Japan) mice, together with a ribonucleoprotein complex comprising Cas9 protein and a guide RNA targeting the Brca2 stop codon by microinjection. Similarly, a *Gclm*^Hypo^ knock-in mouse strain was established by inserting AID tag to the C-terminus of the glutamate-cysteine ligase modifier subunit (Gclm, NCBI Gene ID: 14630). These mice were crossed with *Rosa26*^CAG-OsTIR^ mice, which express OsTIR and IRES-EGFP under the control of the CAG promoter, located on the *Rosa26* locus^40^. Genotype of these mice was confirmed by PCR using genomic DNA extracted from tail biopsies. Primer sequences are listed in Supplemental Table S7.

### *In vivo* Brca2 depletion

Eight– to twelve-week-old *Brca2*^AID/AID^*Rosa26*^CAG-OsTIR/+^ mice were intraperitoneally (i.p.) injected with 5-Ph-IAA (15 mg/kg; A1440350, Ambeed). After 12 hours, Lineage^−^Sca-1^+^c-Kit^+^ (LSK) cells were isolated from bone marrow. Control LSK cells were collected from untreated littermates. Total RNA was extracted using the RNeasy Mini Kit (74104, QIAGEN) and subjected to RNA-seq analysis.

To assess the physiological impact of Brca2 depletion, *Brca2*^AID/AID^*Rosa26*^CAG-OsTIR/+^ mice were treated with cisplatin (2 mg/kg, i.p., 4291401A1097, Nippon Kayaku) with or without N-acetyl-L-cysteine (NAC, 50 mg/kg) on day 0. The 5-Ph-IAA (15 mg/kg, i.p., A1440350, Ambeed) was administered daily for 14 days. Age-matched *Brca2*^AID/AID^*Rosa26*^+/+^ mice served as controls. Body weight was monitored daily, and mice exhibiting >30% weight loss were euthanized per humane endpoint criteria.

To assess sensitivity to PARP inhibitor in Brca2-depletion mouse, *Brca2*^AID/AID^*Gclm*^Hypo/Hypo^*Rosa26*^CAG-OsTIR/+^, *Brca2*^+/+^*Gclm*^Hypo/Hypo^*Rosa26*^+/+^ and *Brca2*^AID/AID^*Gclm*^+/+^*Rosa26*^CAG-OsTIR/+^ mice were treated with Olaparib (25, 12.5, 6.25 or 1.25 mg/kg, i.p., A134565, Ambeed) with 5-Ph-IAA (15 mg/kg, i.p., A1440350, Ambeed). Age-matched *Brca2*^AID/AID^*Gclm*^+/+^ *Rosa26*^+/+^ mice served as controls. Body weight was monitored daily, and mice exhibiting >30% weight loss were euthanized per humane endpoint criteria.

### Isolation and analysis of hematopoietic cells from AID-mice

Bone marrow cells were harvested, erythrocytes were lysed using RBC lysis buffer (420301, BioLegend), and non-specific antibody binding was blocked with Mouse BD Fc Block™ (553142, BD Biosciences). Cells were then stained with biotinylated lineage (Lin) antibodies against CD3ε, Gr-1, B220, Ter-119, TCRβ, TCRγδ, CD19, CD11c and CD11b, followed by PE, APC or Brilliant Violet 421-anti-Sca-1 and APC or PE-anti-c-Kit antibodies. Streptavidin–APC or Brilliant Violet 421 (405226, BioLegend) was used for secondary labeling. Flow cytometric analysis was performed using LSRFortessa™ and FlowJo software (BD Biosciences).

For LSK cell isolation, Lin^−^ cells were collected using MojoSort™ streptavidin nanobeads (480015, BioLegend) and MojoSort™ magnetic separation (BioLegend). Then, Lin^−^ cells were stained with Sca-1 and c-Kit antibodies. LSK cells were sorted using FACSAria™ III (BD Biosciences). T and B cells were isolated from splenocytes using biotinylated anti-TCRβ and anti-CD19 antibodies, respectively, followed by MojoSort™ magnetic separation. HSPCs were cultured in F12 medium supplemented with 1% ITS-X, 10 mM HEPES, 1% P/S, 100 ng/mL mouse TPO, 10 ng/mL mouse SCF, and 0.1% PVA. T cells and B cells were cultured in RPMI with 5% FBS, stimulated with anti-CD3ε/CD28 or LPS (L2880, Sigma-Aldrich) and IL-4 (574304, BioLegend), respectively.

### Generation of Lhx2-LSK Cells

Single-cell suspensions from bone marrow of *Brca2*^AID/AID^*Rosa26*^CAG-OsTIR/+^, *Gclm*^Hypo/Hypo^*Rosa26*^CAG-OsTIR/+^, *Brca2*^AID/AID^*Gclm*^Hypo/Hypo^*Rosa26*^CAG-OsTIR/+^ and mice were prepared. The cells were treated with Mouse BD Fc Block™ Reagent (553142, BD Biosciences) to block non-specific antibody binding, followed by PE-conjugated anti-mouse lineage cocktail (Lin; 133303, BioLegend,) consisting of antibodies against CD3ε (clone 145-2C11), Gr-1 (clone RB6-8C5), B220 (clone RA3-6B2), Ter-119 (clone Ter-119) and CD11b (clone M1/70). Simultaneously, the cells were stained with APC-conjugated anti-Sca-1 (clone D7, 108111, BioLegend) and PE/Cy7-conjugated anti-c-Kit (clone 2B8, 105813, BioLegend) antibodies.

As IRES-EGFP is inserted downstream of *OsTIR* on *Rosa26* locus in *Rosa26*^OsTIR^ mice, EGFP^+^Lin^−^Sca-1^+^c-Kit^+^ (LSK) cells were isolated by FACSAria™ III. Forty to fifty thousand LSK cells were isolated and cultured for 2 or 3 days in IMDM (11506-05, Nacalai Tesque) supplemented with 10% fetal bovine serum (FBS; FB-1345/500, biosera, Cholet, France), NEAA (06344-56, Nacalai Tesque), 55 µM 2-ME (21985-023, Gibco), penicillin-streptomycin mixed solution (26253-84, Nacalai Tesque), referred to as IMDM/FBS medium, and with 10 ng/mL mouse IL-3 (213-13, Peprotech, Rocky Hill, NJ), 20 ng/mL human IL-6 (200-06, Peprotech), and 100 ng/mL mouse SCF (250-03, Peprotech).

The retroviral vector, pMY-Lhx2-IRES-EYFP, was transfected into PLAT-E packaging cells using Lipofectamine™ 2000 (11668019, Invitrogen) according to the manufacturer’s instructions. The retrovirus supernatant was collected and used for the transduction of Lhx2 into the LSK cells. The LSK cells were suspended in 1 mL of filtered retrovirus supernatant in the presence of 8 µg/mL polybrene (H9268, Sigma-Aldrich) and transferred into 1 well of a 24-well plate. The plate was then centrifuged at 1,100 x g for 2 hours at 25°C. The cells were recovered and cultured in IMDM/FBS medium containing 20 ng/mL human IL-6 and 100 ng/mL mouse SCF.

Four to six days later, Lhx2-transduced EYFP^+^ LSK cells were isolated by FACSAria™ III. Optical filter sets were installed to detect EGFP and EYFP simultaneously (535AF LP mirror/520DRLP BP filter for EYFP and 510AF LP mirror/502LP BP filter for EGFP). The isolated cells were cultured as above. The flowcytometric data were processed using FlowJo software.

### Chromosomal aberration analysis

Lhx2-LSK cells derived from *Brca2*^AID/AID^*Rosa26*^CAG-OsTIR/+^ and *Brca2*^AID/AID^*Gclm*^Hypo/Hypo^*Rosa26*^CAG-OsTIR/+^ mice in the logarithmic growth phase were used. These cells were cultured with or without 100 nM 5-Ph-IAA for 16 hrs. Thereafter, 200 nM nocodazole was added for 1 ∼ 2 hrs to induce M-phase arrest. Then, the cells were treated with hypotonic solution (75 mM KCl) and incubated for 10 min at 37°C and fixed with Carnoy’s solution (3:1 mixture of methanol and acetic acid). The cells were then dropped onto a microscope slide and stained with Giemsa solution. For observation, 40 M-phase spreads were scored for each sample using a Keyence BZ-X700 microscope and a 100x oil-immersed lens.

### Quantification and Statistical Analysis

All experiments were independently repeated at least three times. Statistical analyses were performed using GraphPad Prism software. Data are presented as mean ± standard deviation unless otherwise noted.

**Table S6:**
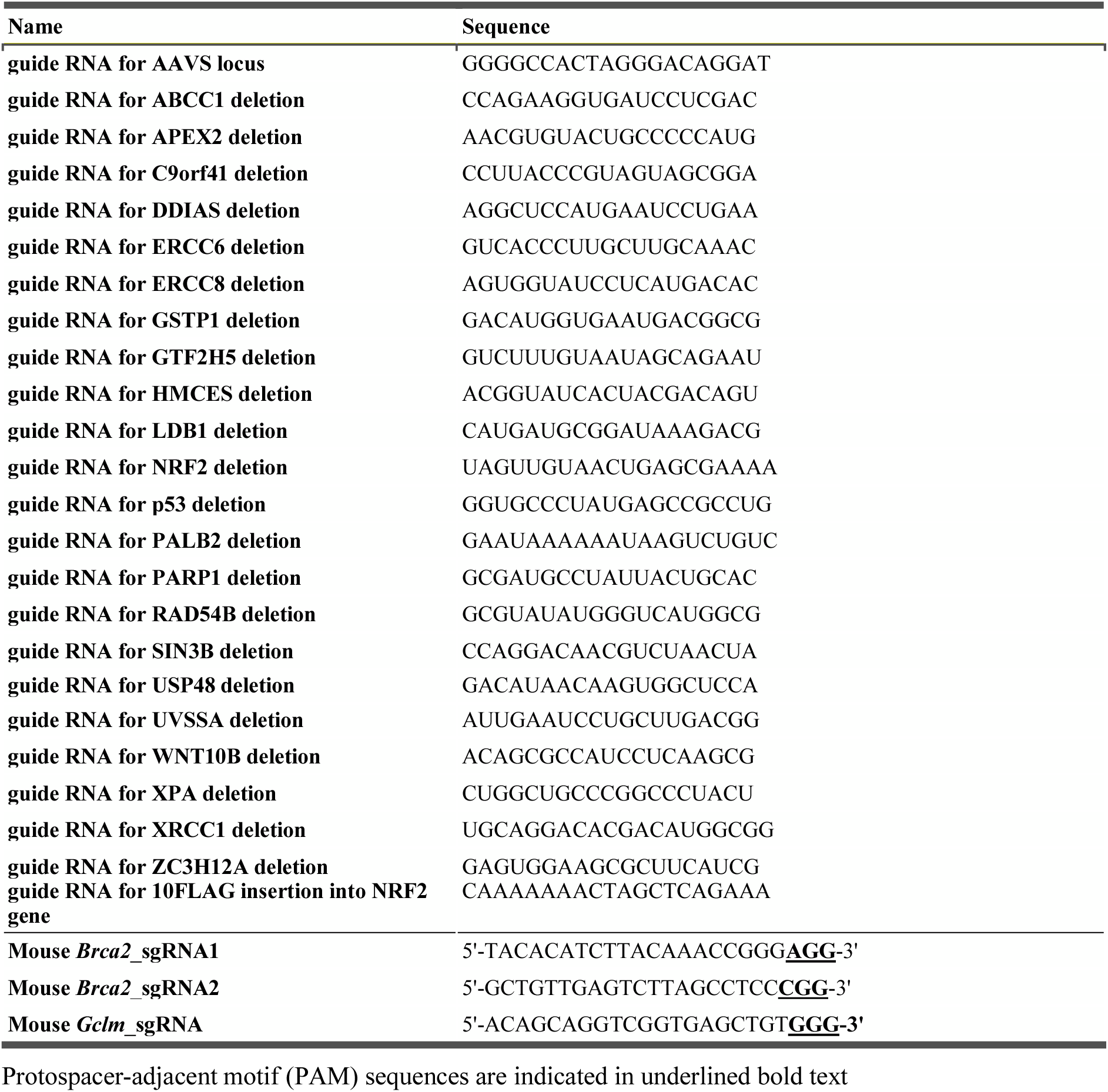
Guide RNA used in this study.

**Table S7:**
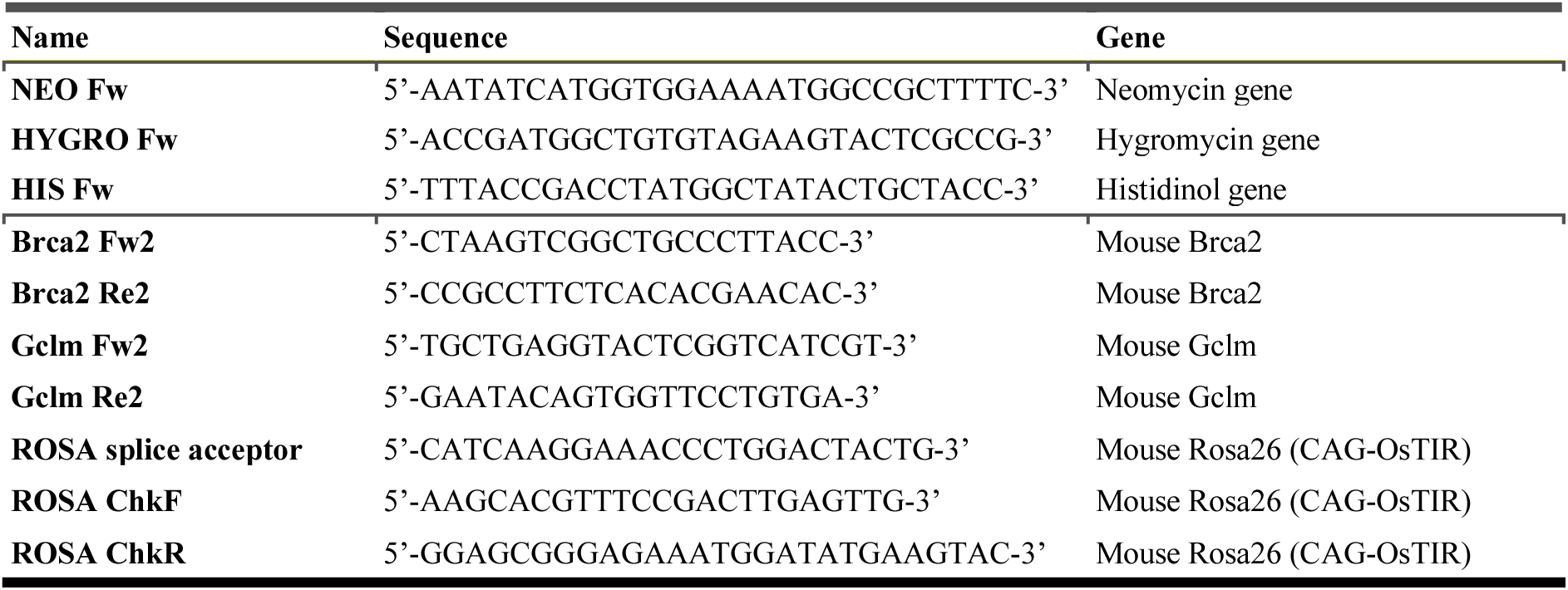
Primers used in this study.

**Table S8:**
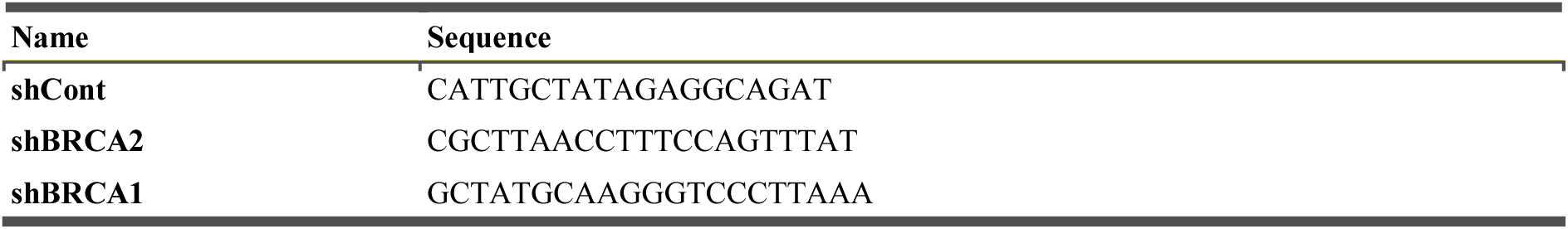
Short hairpin RNA used in this study.

## Supplemental Figure legends

**Figure S1.** Validation of the screen and BRCA2 depletion effects in various cell lines related to Figure 1. (A) Schematic of the mAID knock-in strategy targeting the C-terminus (upstream of the termination codon, T.C.) of *BRCA1*, *BRCA2* and *PALB2*. (B) PCR genotyping of *BRCA1*^AID^, *BRCA2*^AID^, and *PALB2*^AID^ clones using primers targeting the marker gene and 3’ UTR (C) Auxin dose-dependent degradation of BRCA2^AID^ protein (top) and corresponding quantification of RAD51 foci formation 4 h following X-ray-induced DNA damage (bottom). Rad51, DNA, and Cyclin A were colored with green, blue, and red, respectively. Scale bar = 10 µm. (D) Cell proliferation of *BRCA2*^AID^ cells treated with (blue line) or without (gray line) 30 µM auxin. (E) Synthetic lethality screen between *BRCA2* depletion and candidate gene knockouts. Left: experimental scheme, Middle: relative growth ± 30 µM auxin; Right: fold change (auxin+/-) highlighting DNA repair genes (red) and other genes (blue). (F) Identification of genes rescuing BRCA2-deficiency-induced growth defects. Red: DNA repair-related genes; Yellow: other genes (score <10^−4^). (G and H) Growth (G) and survival (H) of *BRCA2*^AID^ cells with *TP53*^+/+^ or *TP53*^−/−^ backgrounds ± 500 µM auxin. (I) Western blots confirming depletion of XRCC1 (left) and NRF2 (right) in *BRCA2*^AID^ cells. (J) H_2_O_2_ dose-response survival curves of 30 µM auxin-treated *BRCA2*^AID^ cells, comparing *XRCC1*^−/−^ (squares) and control (circles). The survival was measured by colony formation assay. (K) Western blot showing depletion of BRCA2 (upper) and BRCA1 (lower) in *wild-type* and *FEN1*^−/−^ cells. (L) Growth curves of *FEN1*^+/+^ and *FEN1*^−/−^ cells transduced with shRNAs targeting control or *BRCA1*. (M) ROS levels in *BRCA2*^AID^ cells with *TP53*^+/+^ or *TP53*^−/−^ following treatment with 500 µM auxin. (N and O) Impact of 100 µM α-tocopherol on ROS levels (N) and growth kinetics (O) in auxin-treated *BRCA2*^AID^ cells. (P) Western blot showing BRCA2 in CAPAN1 (left) and PEO1 (right) cells with (blue circle) or without (white circle) *piggybac*(*pb*)*-BRCA2* expression. (Q and R) Dose-dependent survival of CAPAN1 (left) and PEO1 (right) cells with (squares) or without (circles) expression under treatment with Olaparib (Q) or H_2_O_2_ (R). The survival was assessed by colony formation assay. (S) Images showing ROS (green) and nuclei stained with DAPI (blue) in CAPAN1 cells without *pbBRCA2* (top), with *pbBRCA2* (middle), or treated with N-acetyl-L-cysteine (NAC; bottom). Data are presented as mean ± SD from at least three-independent experiments. Statistical significance was calculated using an unpaired t-test (D, G, H, L, and M–O) or Mann-Whitney test (J and R). \**p* < 0.05, \*\**p* < 0.01, \*\*\**p* < 0.001, \*\*\*\**p* < 0.0001.

**Figure S2.** Depletion of BRCA1 or PALB2 also induces single-strand DNA breaks, related to Figure 2. (A) Western blot of BRCA1 depletion (upper) and PALB2 depletion (lower) in the indicated AID-tagged cells ± 30 µM auxin (6 h). (B) Representative images of the alkaline comet assay in *BRCA1*^AID^ (left) and *PALB2*^AID^ (right) cells with or without 500 µM auxin treatment. Quantification of tail moment following auxin treatment is shown at the bottom. (C) Immunoblot showing p53 stabilization following BRCA2 depletion induced by 500 µM auxin.

**Figure S3.** 8-oxoG blot of breast tumors and normal tissues, related to Figure 2. (A and B) Tumor location and prophylactic resection sites for each patient. (C) Slot blot of 8-oxoG in clinical specimens. Upper panel: normal and tumor tissues from patient#1 with *BRCA* germline variants (*BRCA1* p.Leu63Ter). Lower panel: normal and tumor samples from patient#2 (*BRCA2* p.Tyr2154Ter) and sporadic breast cancer samples. Following DNA quantification, 50 ng of genomic DNA were analyzed per spot. Tumor genotypes and subtypes were analyzed by whole-exome sequencing and clinical diagnosis. The spot to the right of KB20 indicates a failed sample.

**Figure S4.** Correlation between genes upregulated upon BRCA2 depletion and genes accumulating 8-oxoG, related to Figure 2. (A) Overview of the 8-oxoG-seq procedure using anti-8-oxoG antibody with λDNA spike-in as an internal control. (B) Correlation of biological replicates for DMSO– and auxin-treated samples. (C) Venn diagram of 8-oxoG peaks ± auxin treatment. (D–F) Heatmaps and average profiles showing normalized 8-oxoG signal and GC content at common peaks (C), DMSO-only peaks (D), and auxin-only peaks (E), aligned across gene bodies (TSS to TES) with ± 2 kb flanking regions. (G) Genes associated with 8-oxoG peaks (H) Volcano plot of differentially expressed gene upon auxin treatment. Red dots indicate genes harboring 8-oxoG peaks. RNA-seq data is same as in Fig. 5A. (I) Box plots of log_2_(fold-change) in gene expression upon BRCA2 depletion for genes with v.s. without 8-oxoG peaks (Wilcoxon rank-sum, *p* = 7.1×10⁻⁹). (J) Proportions of genes (with/without 8-oxoG peaks) across expression-change categories and standardized Pearson residuals from the chi-square test.

**Figure S5.** Olaparib-induced PARP1 trapping in G1 triggers apoptosis in BRCA2-deficient cells, related to Figure 3. (A) Western blot showing mono/poly (ADP ribose) (PAR) with or without 500 µM auxin treatment for 24 h, combined with 0, 0.1, 1, or 10 µM PARG inhibitor treatment for 6 h. (B) Distribution of Annexin V negative (blue dots) and Annexin V positive (red dots) cells after treatment with 500µM auxin for 72 h (left) ± 10µM Olaparib for 48 h (right). (C) Quantification of the total number of events classified as Annexin V negative or Annexin V positive in the plots shown in (B), under auxin treatment (light blue bars) or auxin plus Olaparib treatment (yellow bars). (D) Flow cytometry analysis showing cell cycle distribution under auxin treatment (left) and auxin plus Olaparib treatment (right) in *BRCA2*^AID^ cells. Each population is indicated by color: early S (light green), late S (green), G_2_ (orange), G_1_ (red), and sub-G_1_ (gray). (E) Time-course of live cell (red line) and Annexin V positive cell (blue line) counts per well over 24 h following treatment with auxin alone (left) or auxin plus Olaparib (right) (F) Flow cytometry analysis showing cell cycle distribution under auxin plus Olaparib treatment in *BRCA2*^AID^ *TP53*^−/−^ cells.

**Figure S6.** BRCA2 is essential for hematopoietic stem cell maintenance and redox homeostasis in *vivo*, related to Figure 4. (A) Schematic diagram illustrating the mAID knock-in strategy upstream of the termination codon in *Brca2* (upper). *OsTIR1* (*F74G*) and GFP are co-expressed from a bicistronic transcript (*IRES-GFP*) driven by the *CAG* promoter inserted into the *Rosa26* locus (bottom). (B) BRCA2-mAID protein remains stable without OsTIR1 expression (left), but is efficiently degraded upon 5-Ph-IAA injection in the presence of OsTIR1 (right). (C) Genotyping of *Brca2*^WT^ and *Brca2*^AID^ by PCR. (D) Evaluation of Mendelian inheritance in offspring from *Brca2*^WT/AID^ mouse crosses. Genotyping of 70 pups showed genotype ratios consistent with expected Mendelian inheritance. (E) Images showing IR-induced RAD51 foci (magenta) and Hoechst-stained nuclei (blue) in thymocyte with or without 5-Ph-IAA treatment (left). RAD51-focus intensities on each nucleus were quantified using imageJ software (right). (F) Body weight of mice expressing OsTIR1 (blue line) or not expressing OsTIR1 (black line) after injection of 5-Ph-IAA and cisplatin. (G) Experimental scheme for 5-Ph-IAA, cisplatin, and N-acetyl-L-cysteine (NAC) injection followed by dissection. (H and I) Relative growth of T cells (H) and B cells (I) in a 5-Ph-IAA concentration-dependent manner (0, 0.001, 0.01, 0.1, and 1 µM). (J and K) CellROX fluorescence intensity indicating ROS levels in T cells (J) and B cells (K) in a 5-Ph-IAA concentration-dependent manner. (L) Images showing Lhx2-LSK (Lin-, Sca-1^+^, and c-Kit^+^) cells after 7-day treatment with IL-6 and SCF (left), and differentiated neutrophil (red arrowhead) and macrophage (green arrowhead) after 7-day treatment with IL-3 and GM-CSF treatment (right), scale bar, 20 µm. (M) Western blot showing BRCA2 expression in Lhx2-LSK after treatment with 100 nM 5-Ph-IAA for 1 day. (N) Images showing RAD51 foci (red) and Hoechst-stained nuclei (blue) in Lhx-LSK cells treated with or without 100 nM 5-Ph-IAA for 1 day, followed by 2 hours of ± 2 Gy irradiation (left). Graph showing quantification of RAD51 foci per cell (right), scale bar, 50 µm. (O) Amounts of various blood cells (WBC, RBC, HGB, and HCT) after 5 consecutive days of treatment with Olaparib (0, 3.125, 6.25, 12.5, or 25 mg/kg) and 5-Ph-IAA (15 mg/kg) in mice with or without OsTIR1 expression. Data are presented as mean ± SD from at least three-independent experiments. Statistical significance was calculated using a chi-squared test (D), an unpaired t-test (N and O) and Mann-Whitney test (E, H-K). \**p* < 0.05, \*\**p* < 0.01, \*\*\**p* < 0.001, \*\*\*\**p* < 0.0001.

**Figure S7.** BRCA2 deficiency creates a glutathione-dependent redox vulnerability, related to Figure 5. (A) Oxygen consumption rate (OCR) profile 6 h post-treatment with 500 µM auxin. (B) Immunoblot of GGT1 expression in *wild-type* and *BRCA2*^AID^ cells ± 500 µM auxin (24 h). Tubulin is a loading control. (C and D) GGT1 activity (FITC intensity) measured by flow cytometry in *BRCA1*^AID^, *BRCA2*^AID^, and *PALB2*^AID^ cells ± 500 µM auxin (24 h). Quantification (C) and representative flow cytometry plots are shown (D). (E) Immunoblot of proteins involved in glutamine/glutathione metabolism (KGA/GAC, GSS, xCT). (F and G) Cell survival of *BRCA1*^AID^ (F) and *PALB2*^AID^ (G) cells with (blue line) or without (gray line) 100 µM auxin treatment in a BSO concentration-dependent manner. The survival was measured by colony formation assay.

**Figure S8.** Combined BRCA2 and GCLM deficiency synergistically drives genomic instability and hematopoietic collapse, related to Figure 6. (A) Schematic diagram illustrating the mAID knock-in strategy upstream of the termination codon in the *Gclm* gene (left). Genotyping of *Gclm*^WT^ and *Gclm*^Hypo^ mice (right). (B and C) Total glutathione concentrations in the liver (B) and erythrocytes (C) isolated from *Gclm*^WT^ and *Gclm*^Hypo^ mice. (D) Western blot showing GCLM and GCLC expression in *Gclm*^Hypo^ Lhx2-LSK cells after treatment with or without 1 µM 5-Ph-IAA for 1 day. (E) Body weight of male (left) and female (right) mice harboring *Gclm*^WT^ and *Gclm*^Hypo^. (F and G) Relative body weight of mice (F) and survival probability (G) following daily IP injections of 5-Ph-IAA (15 mg/kg) for 14 days and a single injection of PHZ (35 mg/kg) on day 0. (H) Amounts of various blood cells (WBC, RBC, HGB, and HCT) after 5 consecutive days of treatment with Olaparib (25 mg/kg) and 5-Ph-IAA (15 mg/kg) in *Brca2*^AID^ mice with or without expression of the hypomorphic (Hypo) form of *Gclm*. Data are presented as mean ± SD from at least three-independent experiments. Statistical significance was calculated using an unpaired t-test (B, C, E, and H). \**p* < 0.05, \*\**p* < 0.01, \*\*\**p* < 0.001, \*\*\*\**p* < 0.0001.

**Figure S9.** Metabolic rewiring toward a pro-oxidant state and systemic exhaustion of redox cofactors upon BRCA2 loss, related to Figure 7. (A) Principal Component Analysis (PCA) of metabolomic data from cells treated with 500 µM auxin (blue dots) or solvent control (gray dots). (B) Hierarchical clustering of metabolites from cells treated with 500 µM auxin (blue circle) or solvent control (white circle). In the heatmap, green/red indicates values below/above the mean, respectively. (C) Relative abundance of the metabolites methionine (Met) S-adenosyl-methionine (SAM), S-adenosyl-l-homocysteine (SAH), homocysteine, cystathionine, and cysteine in *BRCA2*^AID^ cells with (blue bars) or without (gray bars) auxin treatment. (D and E) Relative abundance of the metabolites involved in pentose phosphate pathway (D), and nucleotide synthesis (E) with (blue bars) or without (gray bars) auxin treatment. (F) Growth of *BRCA2*^AID^ cells was assessed under four different culture conditions, including glucose, galactose, glucose with auxin, and galactose with auxin. The number of live cells (light blue bars) and dead cells (yellow bars) are shown. Data are presented as mean ± SD from at least three-independent experiments. Statistical significance was calculated using an unpaired t-test (C–F). \**p* < 0.05, \*\**p* < 0.01, \*\*\**p* < 0.001, **** *p* <0.0001.

## Notes

### Competing Interest Statement

The authors have declared no competing interest.

