## Supplementary Figure for "BRCA2 loss triggers a downward spiral of genomic instability via ROS-dependent metabolic collapse"

Hirai,H. *et al*, Figure S1

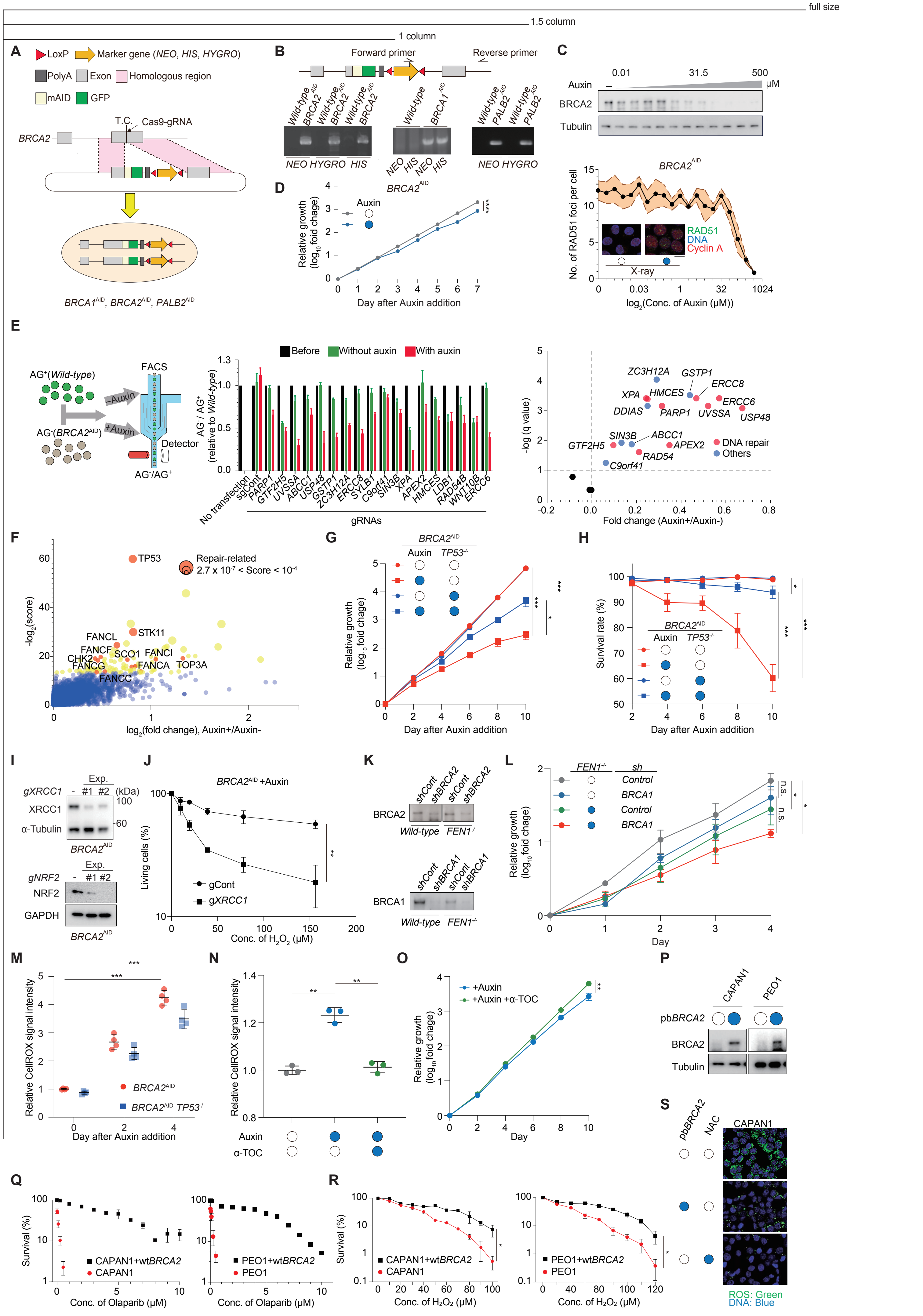

full size

1.5 column

1 column

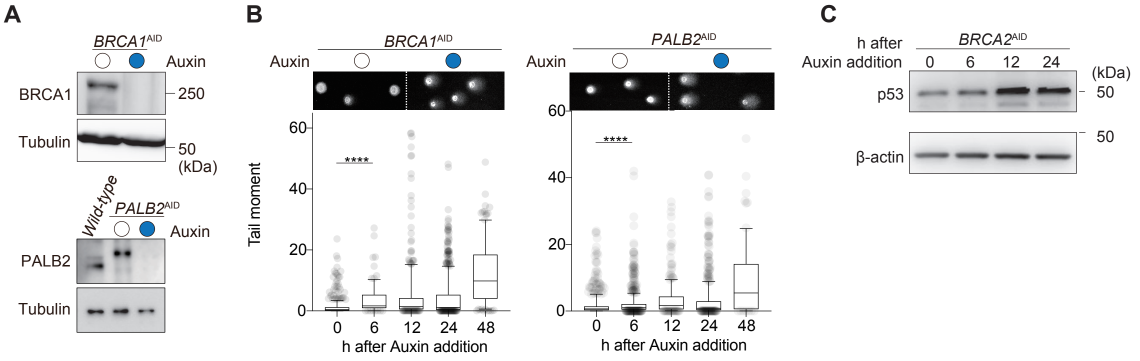

full size

1.5 column

1 column

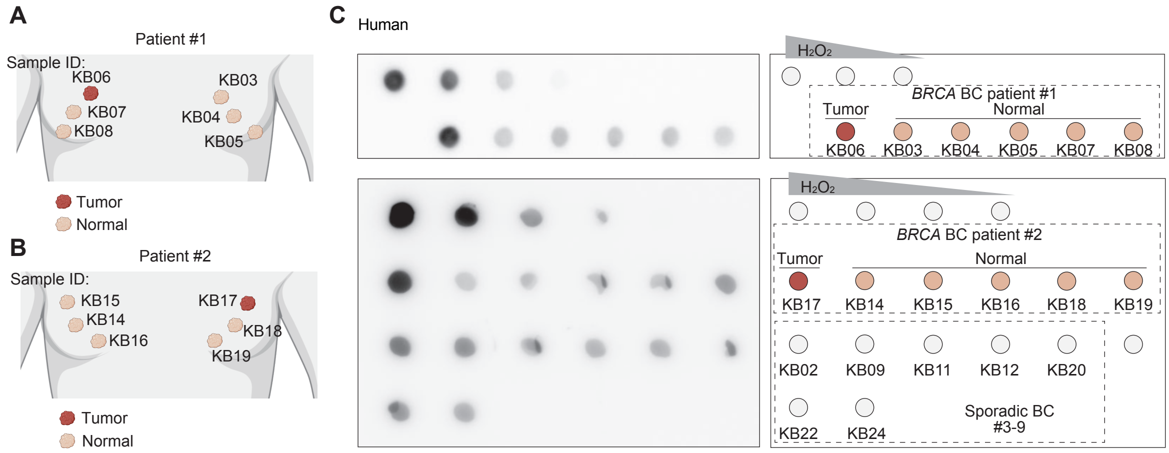

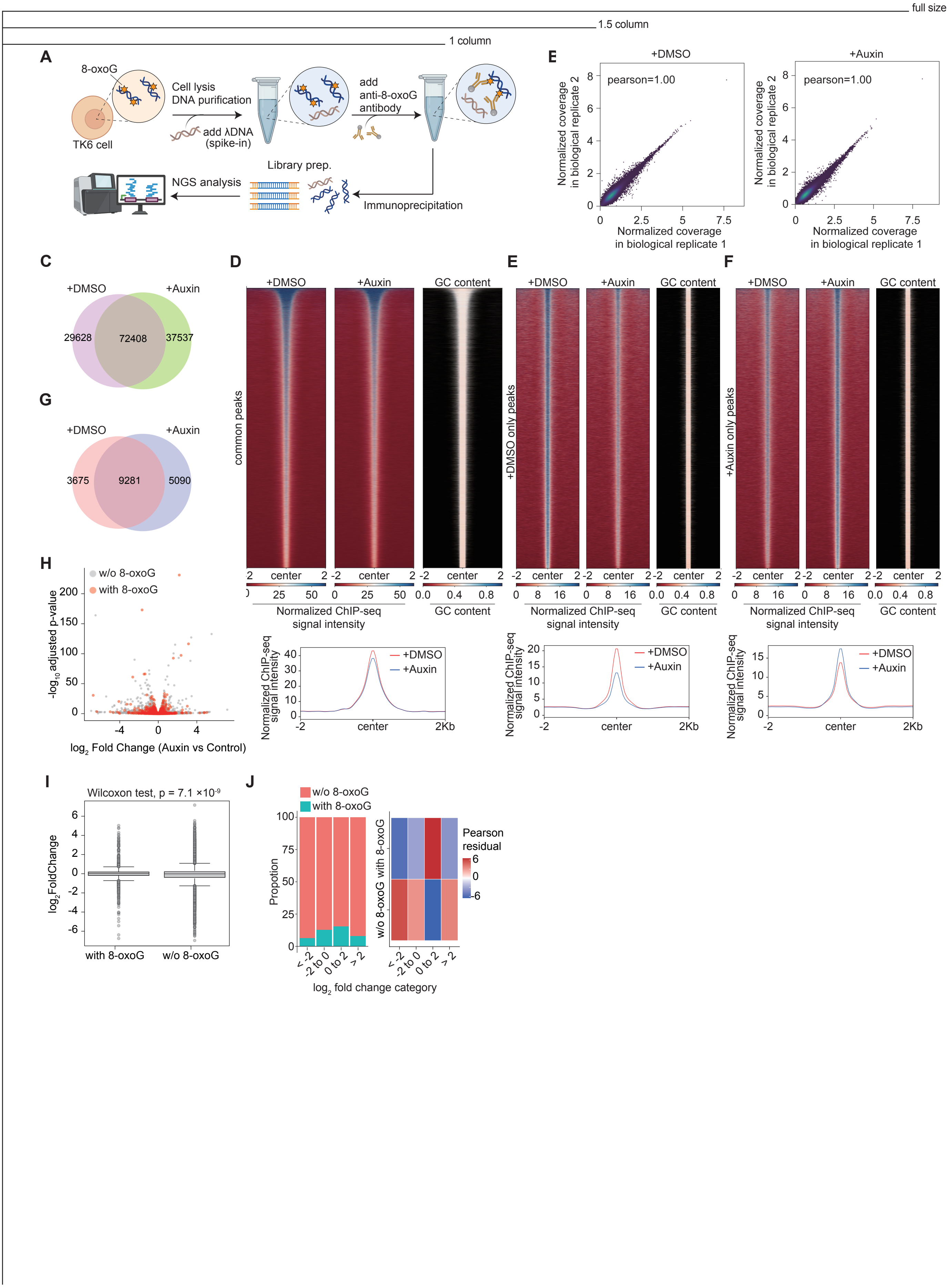
full size
1.5 column
1 column
A
8-oxoG
TK6 cell
Cell lysis
DNA purification
add λDNA
(spike-in)
add
anti-8-oxoG
antibody
Library prep.
NGS analysis
Immunoprecipitation
E
+DMSO
+Auxin
pearson=1.00
Normalized coverage
in biological replicate 2
Normalized coverage
in biological replicate 1
C
+DMSO
+Auxin
29628
72408
37537
G
3675
9281
5090
D
common peaks
GC content
center
Normalized ChIP-seq
signal intensity
E
+DMSO only peaks
F
+Auxin only peaks
H
w/o 8-oxoG
with 8-oxoG
-log10 adjusted p-value
log2 Fold Change (Auxin vs Control)
Normalized ChIP-seq
signal intensity
center
2Kb
I
Wilcoxon test, p = 7.1 ×10-9
log2FoldChange
with 8-oxoG
w/o 8-oxoG
J
w/o 8-oxoG
with 8-oxoG
Propotion
Pearson
residual
log2 fold change category
< -2
-2 to 0
0 to 2
> 2

full size
1.5 column
1 column

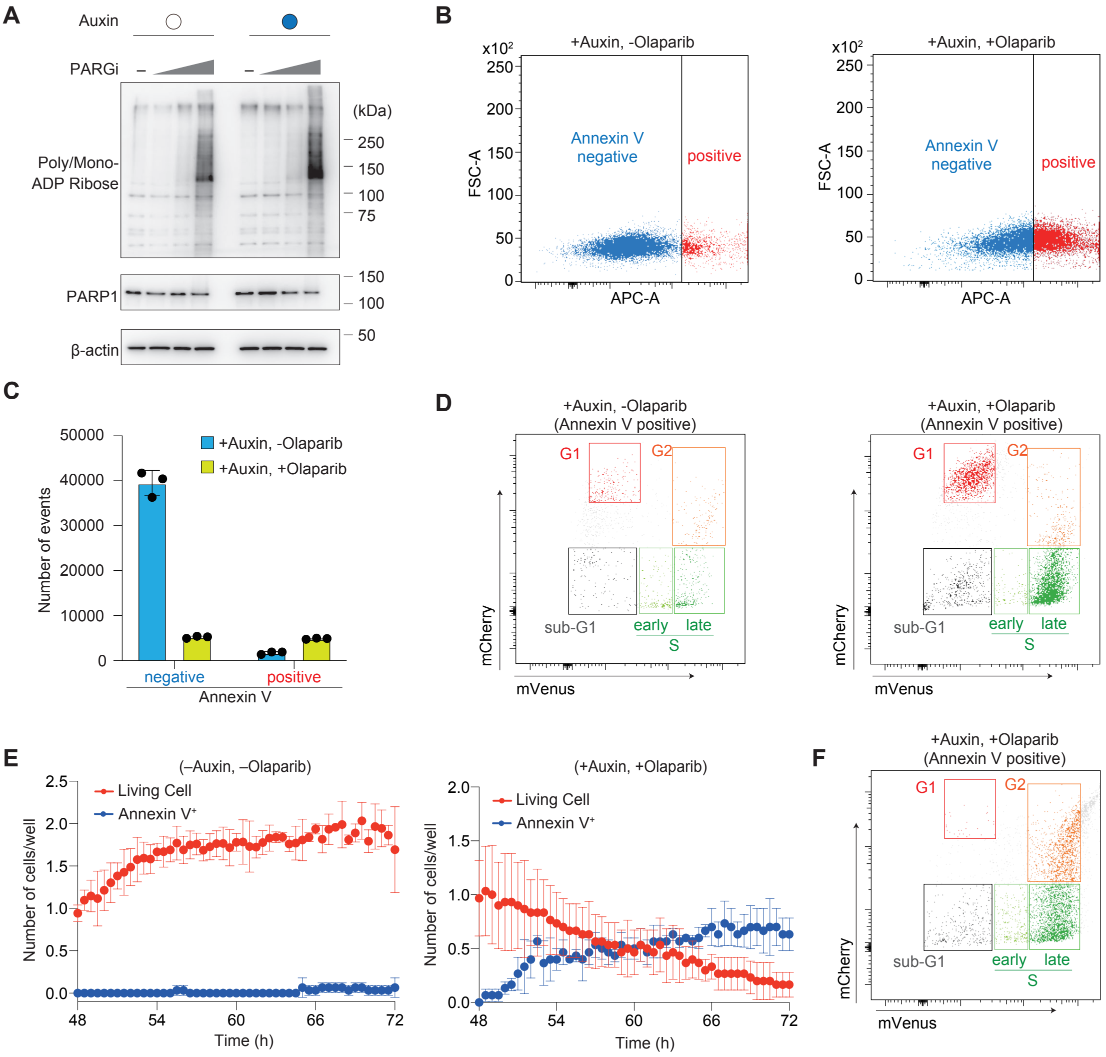

full size

1.5 column

1 column

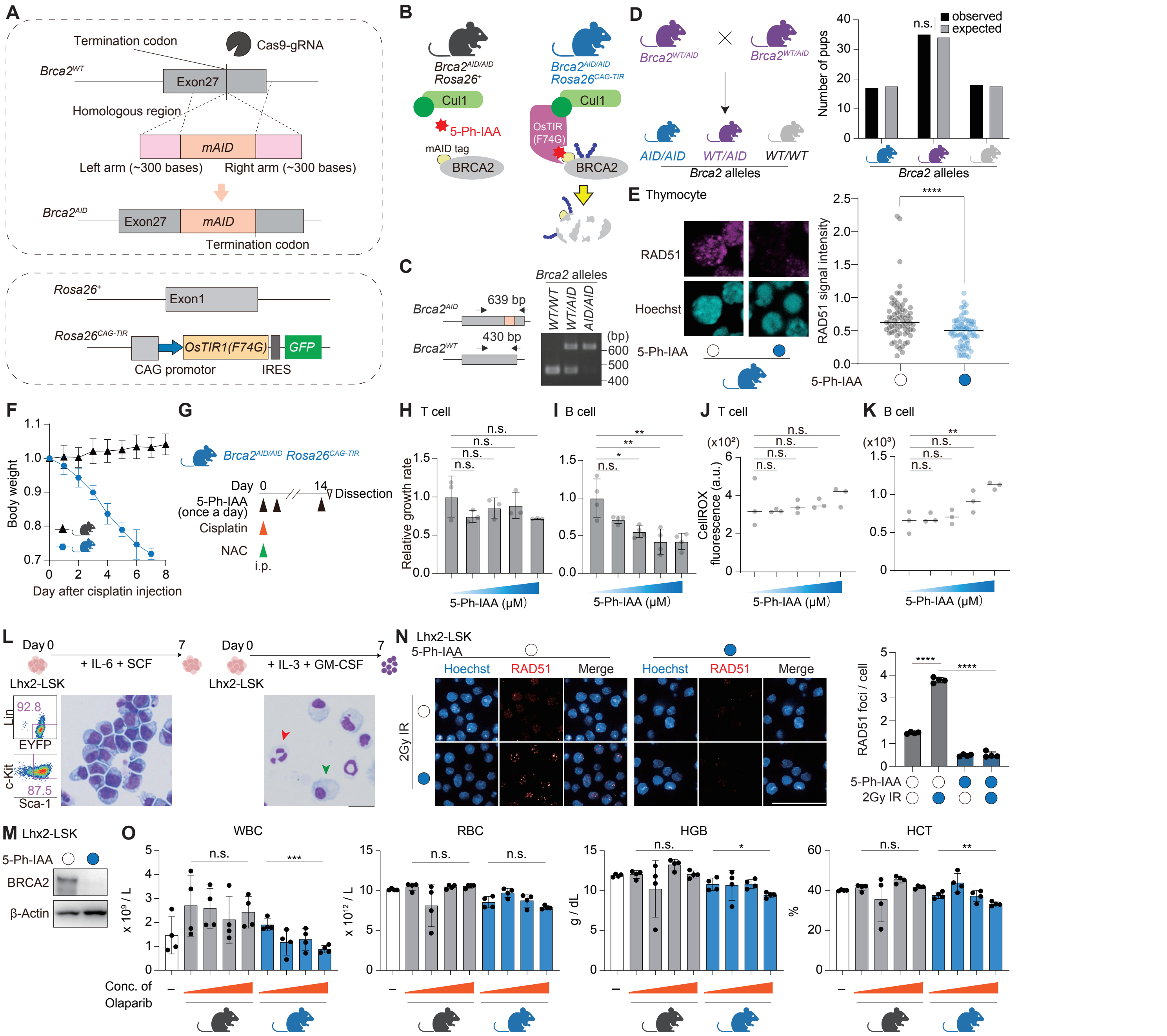

full size

1.5 column

1 column

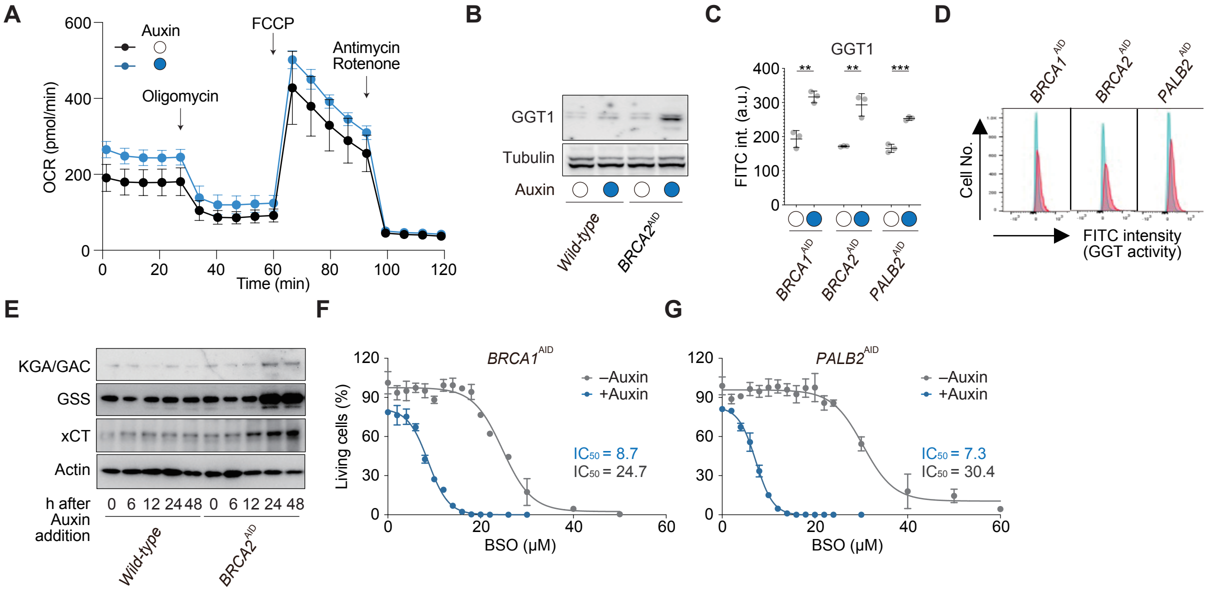

full size

1.5 column

1 column

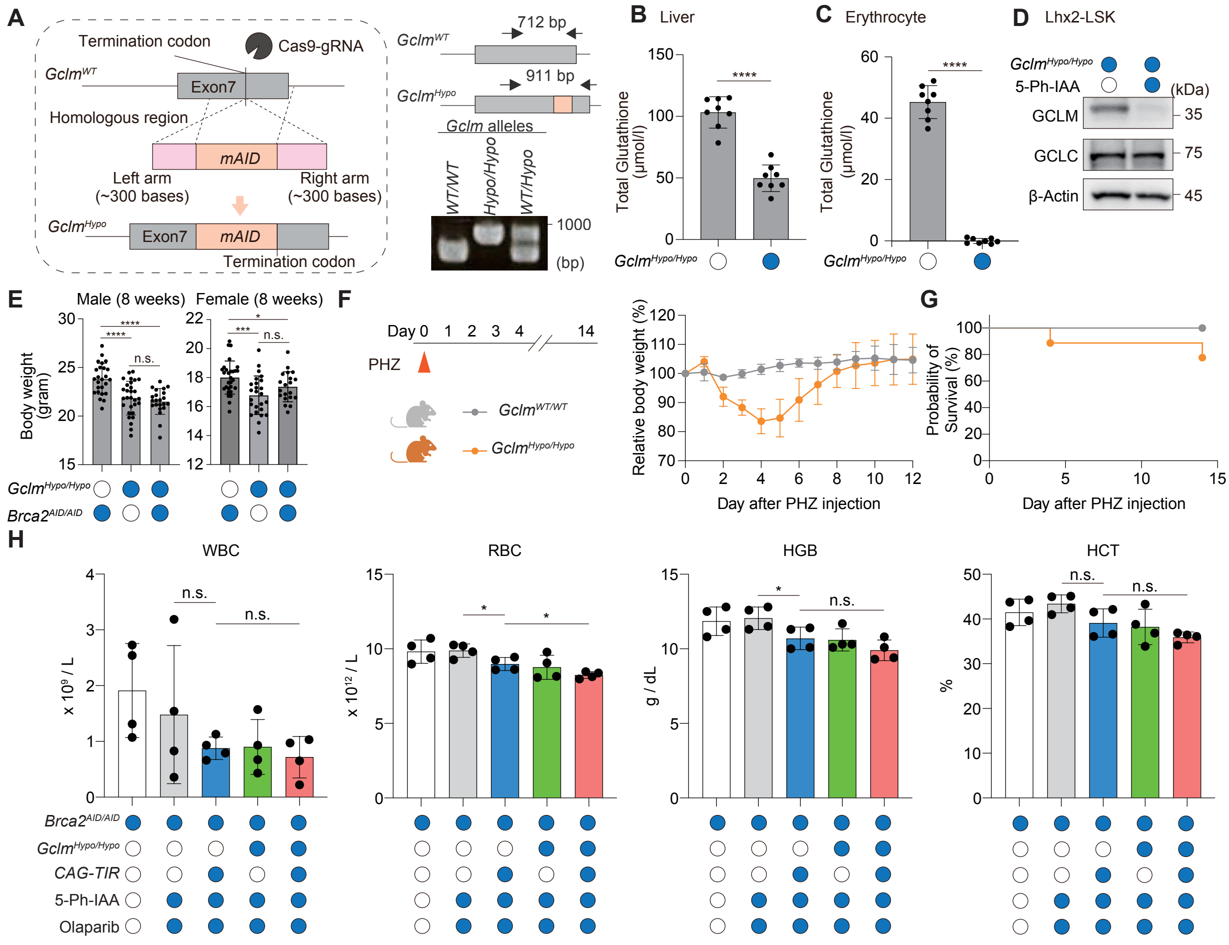

full size

1.5 column

1 column

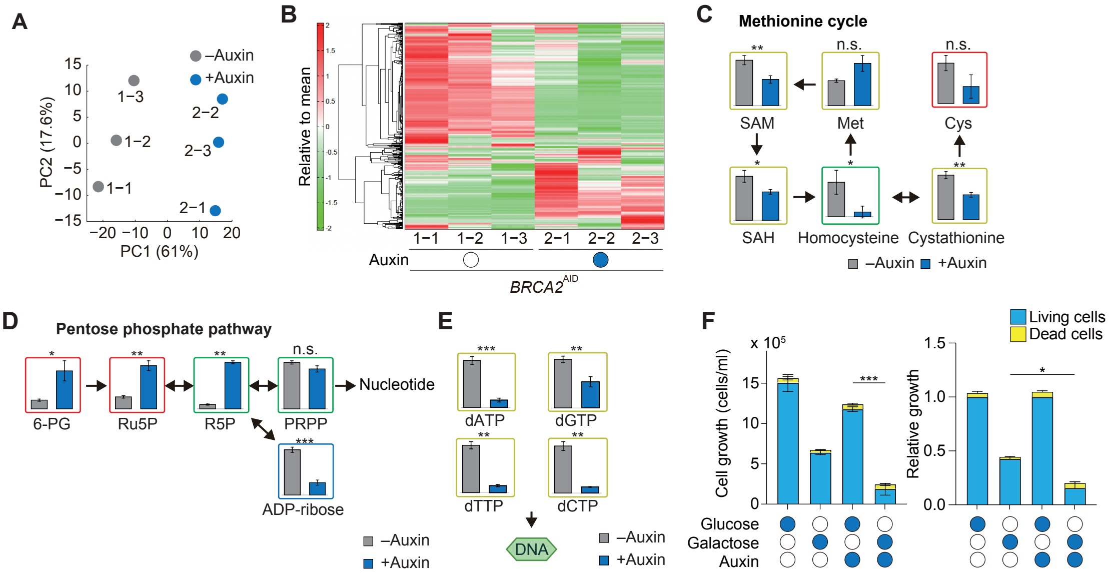
